# Chromosome-scale genome assemblies of five different *Brassica oleracea* morphotypes provide insights in intraspecific diversification

**DOI:** 10.1101/2022.10.27.514037

**Authors:** Chengcheng Cai, Johan Bucher, Richard Finkers, Guusje Bonnema

## Abstract

*Brassica oleracea* is an economically important vegetable and fodder crop species that includes many morphotypes exhibiting enormous phenotypic variations. Previously, a pan-genome study based on short reads mapping approach has shown extensive structural variations between *B. oleracea* morphotypes. Here, to capture more complete genome sequences of *B. oleracea*, we report new chromosome-scale genome assemblies for five different morphotypes, namely broccoli, cauliflower, kale, kohlrabi and white cabbage, which were created by combining long-read sequencing data and Bionano DLS optical maps. The five assemblies are the most continuous and complete *B. oleracea* genomes to date (contig N50 > 10 Mb). Comparative analysis revealed both highly syntenic relationships and extensive structural variants among the five genomes. Dispensable and specific gene clusters accounted for ~38.19% of total gene clusters based on a pan-genome analysis including our five newly assembled genomes and four previously reported genomes. Using the pan-genome of *B. oleracea* and *B. rapa*, we revealed their different evolutionary dynamics of LTR-RTs. Furthermore, we inferred the ancestral genome of *B. oleracea* and the common ancestral genome of *B. oleracea* and *B. rapa* via a pan-genome approach. We observed faster WGT-derived gene loss in *B. rapa* than in *B. oleracea* before intraspecific diversification. We also revealed continuing gene loss bias during intraspecific diversification of the two species and a strong bias towards losing only one copy among the three paralogous genes. This study provides valuable genomic resources for *B. oleracea* improvement and insights towards understanding genome evolution during the intraspecific diversification of *B. oleracea* and *B. rapa*.

## Introduction

Long-read sequencing and long-range scaffolding technologies have significantly improved the quality of genome assemblies. Pacific Biosciences (PacBio) and Oxford Nanopore Technologies (ONT) are two commercial high-throughput long-read sequencing platforms that can generate DNA fragments ranging from kilobases to megabases (Rousseau-Gueutin et al., 2020). Although the raw reads of these technologies have error rates of up to 15%, the accuracy of assembled sequences can reach as high as 99.999% via error correction strategies (Belser et al., 2018; Jiao et al., 2017; Jiao and Schneeberger, 2017). More importantly, long-read sequencing data enables genome assemblies with high contiguity and completeness whereas short-read data (Illumina, Roche 454) often results in highly fragmented assemblies as it is not suitable to span repetitive regions. The last few years have seen highly continuous genomes assembled by long reads for a wide range of species (Belser et al., 2018; Deschamps et al., 2018; Ou et al., 2020; Schmidt et al., 2017). Even though long-read sequencing technologies have enabled highly continuous assemblies, they are solely not sufficient to completely assemble complex eukaryotic genomes, especially those of plants with high levels of repeat sequences and large genome size. The resulting contigs from sequencing reads require further scaffolding to eventually achieve chromosome-scale assemblies. One such long-range scaffolding technology is Bionano Genomics optical mapping with its new Direct Label and Stain (DLS) technology. This technology uses Direct Labeling Enzyme 1 (DLE-1) to attach a single fluorophore to specific sequence motifs to produce fingerprints of DNA fragments, thus not damaging DNA molecules at specific sites (Deschamps et al., 2018). The DLS-labeled molecules are much longer than those labelled via the endonuclease approach, with the longest ones becoming larger than 2 Mbp. Chromosome-scale assemblies can often be generated in single maps by using DLS molecules (Formenti et al., 2019).

*Brassica oleracea* (CC, 2n=18) is an economically important vegetable and fodder crop species cultivated worldwide. It consists of many morphotypes which exhibit enormous phenotypic variations, such as the leafy heading morphotype *var. capitata* (cabbage), the typical curd morphotypes with large arrested inflorescences, including *var. botrytis* (cauliflower), *var. italica* (broccoli), the kohlrabi’s with their tuberous stems (*var. gongylodes*), the kales (*var. acephala*) with different leaf types and etc (Bonnema et al., 2011; Cai et al., 2022a; Dias, 2012; Kole and Henry, 2010). Despite this enormous diversity, *B. oleracea* truly remains one species and morphotypes can be easily interbred. *B. oleracea* is an important diploid member of the “triangle of U” model, which includes the other two diploid species, *Brassica rapa* (AA, 2n=20) and *Brassica nigra* (BB, 2n=16), and three allotetraploid species generated through pairwise crosses between the diploids, *Brassica juncea* (AABB, 2n=36), *Brassica napus* (AACC, 2n=38) and *Brassica carinata* (BBCC, 2n=34) (Nagaharu, 1935). Similar to *B. oleracea, B. rapa, B. juncea* and *B. napus* all include many diverse morphotypes showing extreme phenotypes.

To date, several genome sequences of *B. oleracea* have been created either using short-read or long-read sequencing technology. Two *B. oleracea* reference genomes were firstly released in 2014, one from cabbage line 02-12 and the other from the doubled haploid *B. oleracea* annual kale-like type TO1000DH (Liu et al., 2014b; Parkin et al., 2014a). Both these genome sequences were generated on the basis of Sanger sequencing and deep short-read sequencing data, and anchored to pseudo-chromosomes by genetic maps. These two genomes have been used as the references for many years for comparative and functional genomics studies in *B. oleracea* crops (Cheng et al., 2016b; Golicz et al., 2016b; Zhang et al., 2015). Recently, several other reference genomes, which were primarily created by combining high-coverage long-read sequencing data with long-range scaffolding information, such as optical maps and/or Hi-C, have become available. This includes a re-assembly of cabbage line 02-12 (Cai et al., 2020), a new broccoli line HDEM (Belser et al., 2018), two new cabbage lines (OX-heart and D134) (Guo et al., 2020; Lv et al., 2020) and two new cauliflower lines (Korso and C-8) (Guo et al., 2020; Sun et al., 2019). Increasing numbers of pan-genome studies based on *de novo* assembly approaches in multiple crops, such as rice (Qin et al., 2021; Zhao et al., 2018), soybean (Liu et al., 2020), maize (Hufford et al., 2021) and rapeseed (Song et al., 2020), have shown that structural variations widely exist within a species. Also in *B. oleracea*, a pan-genome study based on a reference-guided approach using short-read sequencing technology shows such high level of variations between different morphotypes, with nearly 20% of genes affected by presence/absence variation (Golicz et al., 2016a). These studies illustrate that a single reference genome is not sufficient to cover the genome sequences of a species. For these reasons, it is necessary to generate more *B. oleracea* reference genomes from highly diverse morphotypes, especially high quality sequences, to fully resolve structural variations that affect gene function and expression and influence agriculturally important traits.

Polyploidization is prevalent and recurrent in the plant kingdom and plays a crucial role in speciation and species diversification (Cai et al., 2021; Cheng et al., 2014; Zhang et al., 2019). *Brassica* species are ideal models for polyploidy and evolutionary studies in plants partly because they have experienced the common Brassiceae-specific whole genome triplication (WGT) event (Liu et al., 2014a; Parkin et al., 2014b; Wang et al., 2011). This event has been confirmed by extensive comparative analyses between genome sequences of *Brassica* species and *Arabidopsis thaliana* (Cheng et al., 2014; Liu et al., 2014a; Wang et al., 2011). In *B. rapa*, the three subgenomes: least fractionated (LF), medium fractionated 1 (MF1) and most fractionated 2 (MF2), were demonstrated to have been evolved from a common translocation Proto-Calepineae Karyotype (tPCK) ancestral diploid genome (Cheng et al., 2013; Cheng et al., 2014). The extant genome structures of diploid *Brassica* species were shaped during the rediploidization process that followed the WGT, which involves extensive gene fractionation, genomic reshuffling and chromosome reduction (Cheng et al., 2014). To illustrate the WGT process from a genome evolution perspective in *Brassica* plants, Cheng et al proposed a “two-step theory” that suggests a tetraploidization event between the tPCK genomes of MF1 and MF2, followed by fractionation and a subsequent hybridization event between the diploidized genome and a third tPCK genome LF (Cai et al., 2021; Cheng et al., 2012a; Cheng et al., 2014). This theory also explains the subgenome dominance observed in *Brassica* plants. Like Brassica’s, many other species including maize, wheat, cotton, grasses, and *A. thaliana* (Akama et al., 2014; Buggs et al., 2010; Cheng et al., 2016a; Cheng et al., 2012a; Li et al., 2014; Murat et al., 2014; Pont et al., 2013; Renny-Byfield et al., 2015; Schnable et al., 2011; Senchina et al., 2003; Wang et al., 2006) display subgenome dominance, with one subgenome retaining more genes, contributing more highly expressed genes and accumulating fewer non-synonymous mutations than other subgenomes (Cheng et al., 2014).

As a pervasive source of genetic change, gene loss is prevalent in all life kingdoms and has great potential to result in adaptive phenotypic diversity (Albalat and Cañestro, 2016; De Smet et al., 2013). It has been found that gene loss is coupled with extensive polyploidy events that create redundant genes, the loss of which usually does not result in apparent functional consequences (Albalat and Cañestro, 2016). In addition to biased gene loss between different subgenomes following polyploidization, gene loss is also biased towards gene function (Albalat and Cañestro, 2016). In *B. rapa* and *B. oleracea*, genes involved in the response to phytohormone signalling were found to be significantly over-retained during gene loss after WGT (Cheng et al., 2014; Wang et al., 2011). Moreover, a functional bias of gene loss can often be observed in species that suffer relaxation of a given biological or environmental constraint. This functionally biased gene loss is caused by the ‘co-elimination’ of genes that are functionally linked in distinct pathways or complexes associated with relaxed constraint (Albalat and Cañestro, 2016; Aravind et al., 2000; Koonin et al., 2004). In a recent research, Cai et al (Cai et al., 2021) inferred the *B. rapa* ancestral genome using a pan-genome based approach, which provides an essential reference to investigate gene loss during intraspecific diversification. They further illustrated the impacts of WGT event and subgenome dominance on intraspecific diversification of *B. rapa*. *B. oleracea* and *B. rapa* are sister species and their divergence occurred at about 4.6 Mya (Cheng et al., 2017). Due to the lack of high quality reference genome sequences, individual genome evolution regarding gene loss during intraspecific diversification of *B. oleracea* is still unexplored. In addition, studies focussing on comparative genome evolution between sister species after speciation from their common ancestor are sparse.

Here, we release chromosome-scale genome assemblies of five *B. oleracea* morphotypes, including broccoli, cauliflower, kale, kohlrabi and white cabbage, all of which were generated by integrating long reads, optical mapping molecules (BioNano Genomics DLS technology) and Illumina short reads. The final assemblies showed extremely high contiguity, with contigs having N50 values between 11.4 Mb and 16.3 Mb and scaffolds having N50 values between 30.5 Mb and 34.1 Mb. Comparative analysis among the five new assemblies demonstrates high degrees of synteny among *B. oleracea* genomes as well as extensive structural variations. Together with four previously published high-quality genomes, we revealed the composition and features of a *B. oleracea* pan-genome via a *de novo* assembly approach. Additionally, we investigated intact LTR-RTs in the pan-genome of *B. rapa* and *B. oleracea*, and compared their evolutionary dynamics. Furthermore, using a pan-genome approach, we studied the impacts of WGT event and subgenome dominance on intraspecific diversification of *B. oleracea*. We also compared evolutionary patterns regarding biased WGT-derived gene loss between *B. rapa* and *B. oleracea* after the speciation from their common ancestor. Together, our work provides valuable resources for genomic-assisted breeding of *B. oleracea* and sheds lights on understanding intraspecific diversification of *B. oleracea* and *B. rapa*.

## Results

### Contig assembly of five *B. oleracea* morphotypes

Five *B. oleracea* accessions (DH lines) representing five different morphotypes, broccoli, cauliflower, kale, kohlrabi and white cabbage, were selected for genome sequencing. Libraries generated from HMW DNA from fresh leaf tissues were used as input to generate sequences for the five genomes on Oxford Nanopore GridION platform. We produced 1.5 to 7.3 million raw ONT long reads for the five morphotypes, totalling 22.5 to 42.7 Gb of data, with N50 values ranging from 13.04 Kb to 30.22 Kb (Table S1). Assuming a 630 Mb *B. oleracea* genome size (Liu et al., 2014a), these nanopore long reads represented 36~68-fold coverage, and sequences longer than 50 Kb amounted to as high as 9.1~11.6-fold coverage. Besides Nanopore long-reads, we also produced 4.2-8.3 Gb (6.7-13.2-fold coverage) PacBio sequences for each morphotype, with N50 values ranging from 17.5 Kb to 20.9 Kb (Table S2). In addition, more than 126.25Gb Illumina reads were generated for each morphotype, covering more than 200-fold coverage of the five genomes (Table S3).

The ONT reads were assembled using SMARTdenovo (Liu et al., 2021b) and the resulting five assemblies all showed high contiguity. They consisted of 315-426 total contigs and featured N50 values from 6.3 to 13.1 Mb (Table S4-S8). Our broccoli raw assembly had a contig N50 size of 9.3 Mb, which is similar to the value of the final HDEM assembly (9.5 Mb, most contiguous released *B. oleracea* reference genome so far) (Belser et al., 2018). The total contig size ranged from 536.6 to 562.7 Mb and the largest contig varied from 20.1 to 35.7 Mb. Due to the absence of an error correction stage in the algorithm of SMARTdenovo (Liu et al., 2021b), the consensus sequences required further polishing to improve base accuracy. We polished raw contigs using both Nanopore and Illumina reads by running two rounds of Racon (Vaser et al., 2017), followed by three rounds of Pilon (Walker et al., 2014). Generally, the polishing process slightly increased N50 values but greatly improved complete BUSCO values (Table S4-S8). The complete BUSCO values were 82.2%-86.4% for raw contigs in the five assemblies. The scores increased to 87.6%-90.4% after Racon polishing and to greater than 97% after Pilon polishing. To further evaluate base accuracy of our assemblies, we mapped Illumina reads to the corresponding genome to identify genomic variations. In total, we identified 3,085,054-3,593,799 variations (SNPs and small InDels) in each of five raw assemblies, however, after the polishing process, only 44,806-57,274 variations were identified, indicating remarkable base quality improvement. The polished assemblies reached high QVs at around 40 and high identities at around 99.99% (see Methods) (Table S9).

### Bionano DLS genome maps generation and hybrid scaffolding

The combination of Bionano Saphyr system and DLS technology yielded 2,767,569-20,443,958 DNA molecules with lengths longer than 20 Kb for each of the five genomes. After filtering out molecules smaller than 150 Kb and molecules with < 9 labeling sites, a total of 96.47-155.61 Gb Bionano molecules, with N50 values ranging from 241.84 to 381.83 Kb, were assembled into genome maps. For each genome, the final *de novo* assembly yielded only 63-93 maps, with N50 values reaching 29.02-33.77 Mb. The resulting total genome map length was 575.94-645.47 Mb, with the largest Bionano map being 47.51-51.00 Mb (Table S10).

To further improve genome assemblies, contigs for each genome were scaffolded with corresponding DLS maps. The resulting hybrid scaffolds showed strong improvements in contiguity, compared to the ONT contig assemblies alone. More than 97.5% of the original ONT contig sequences were anchored to only ~40 scaffolds in each genome, totalling 541.31-557.60 Mb. Scaffold N50 values for the five hybrid assemblies were ~30 Mb and the largest sequence was longer than 46 Mb (Table S11). After resolving 13-bp gaps and gap-filling, we generated the final assemblies for the five genomes, which comprised only 150-249 scaffolds and had contig N50s between 11.43 and 16.28 Mb (Table 1). The remaining gaps accounted for only 1.04-1.49% of the total scaffold size. The long terminal repeat (LTR) assembly index (LAI) (Ou et al., 2018) was 15.56-16.83 in each of the five genomes, indicating high quality of the genome assemblies. We also evaluated the quality by mapping mRNA-seq reads to the corresponding assembly. Up to 98% of mRNA-seq reads can be aligned to the genomes, among which ~94% reads were aligned concordantly exactly one time (Table S23). Together, these results illustrated the high quality of the five *B. oleracea* assemblies. To construct chromosome-level pseudomolecules, we used a homolog-based approach and mapped super-scaffolds to the HDEM reference genome (Belser et al., 2018; Jiao and Schneeberger, 2020). More than 94% scaffold sequences were anchored to the nine pseudochromosomes (Table 1).

**Table 1.** Statistics of final genome assemblies.

|  | Broccoli | Cauliflower | Kale | Kohlrabi | White Cabbage |
| --- | --- | --- | --- | --- | --- |
| **Contigs** |  |  |  |  |  |
| Total length (Mb) | 552.24 | 540.33 | 560.01 | 550.61 | 557.03 |
| Total number | 316 | 270 | 364 | 283 | 316 |
| N50 (Mb) | 14.46 | 11.43 | 16.28 | 11.74 | 12.17 |
| Longest (Mb) | 38.59 | 23.43 | 46.44 | 28.36 | 43.79 |
| **Scaffolds** |  |  |  |  |  |
| Total length (Mb) | 558.04 | 547.89 | 568.26 | 557.26 | 565.47 |
| Total number | 226 | 150 | 249 | 162 | 177 |
| N50 (Mb) | 30.57 | 34.06 | 30.34 | 30.5 | 31.59 |
| L50 | 8 | 7 | 8 | 8 | 8 |
| N90 (Mb) | 12.41 | 13.93 | 13.21 | 11.74 | 12.82 |
| L90 | 18 | 16 | 18 | 18 | 17 |
| Longest (Mb) | 47 | 47.41 | 48.64 | 46 | 48.23 |
| Gap size (Mb) | 5.8 | 7.56 | 8.25 | 6.65 | 8.44 |
| GC | 36.72 | 36.65 | 36.71 | 36.71 | 36.74 |
| LAI | 15.56 | 16.83 | 16.07 | 16.83 | 16.31 |
| Chr. length (Mb) | 526.03 | 524.69 | 535.23 | 531.79 | 533.55 |
| Anchor ratio | 94.26% | 95.77% | 94.19% | 95.43% | 94.36% |

### Genome annotation and comparative genomics

We annotated repetitive elements (TEs) for the five genomes using the EDTA pipeline (Ou et al., 2019). Approximately 51.92-53.74% (280.54-299.05 Mb) of the assembled sequences of each genome were composed of TEs (Fig. 1A and 1B, Table S12-S16), similar to both the JZS v2 and HDEM genomes (Belser et al., 2018; Cai et al., 2020). The most abundant TEs were the LTR-RTs, representing 26.13-28.97% (141.21-161.35 Mb) sequences in each of the five genomes. Using an integrated strategy combining *ab initio*, homolog-based and transcript-based prediction, we identified a total of 60,644-61,995 protein-coding genes in each of the five genomes (Table S17). Nearly 99% (98.16-99.25%) of the genes were located on chromosomes in our five assemblies. BUSCO assessment based on 1,440 conserved plant genes showed that 96.80-97.80% complete genes were present in each genome. Gene functional annotation demonstrated that more than 97.30% of the annotated genes in each genome are supported by homology to known proteins or functional domains in other species (Table S18). Taken together, these results showed the near-complete gene models in our five assemblies.

**Fig. 1.**
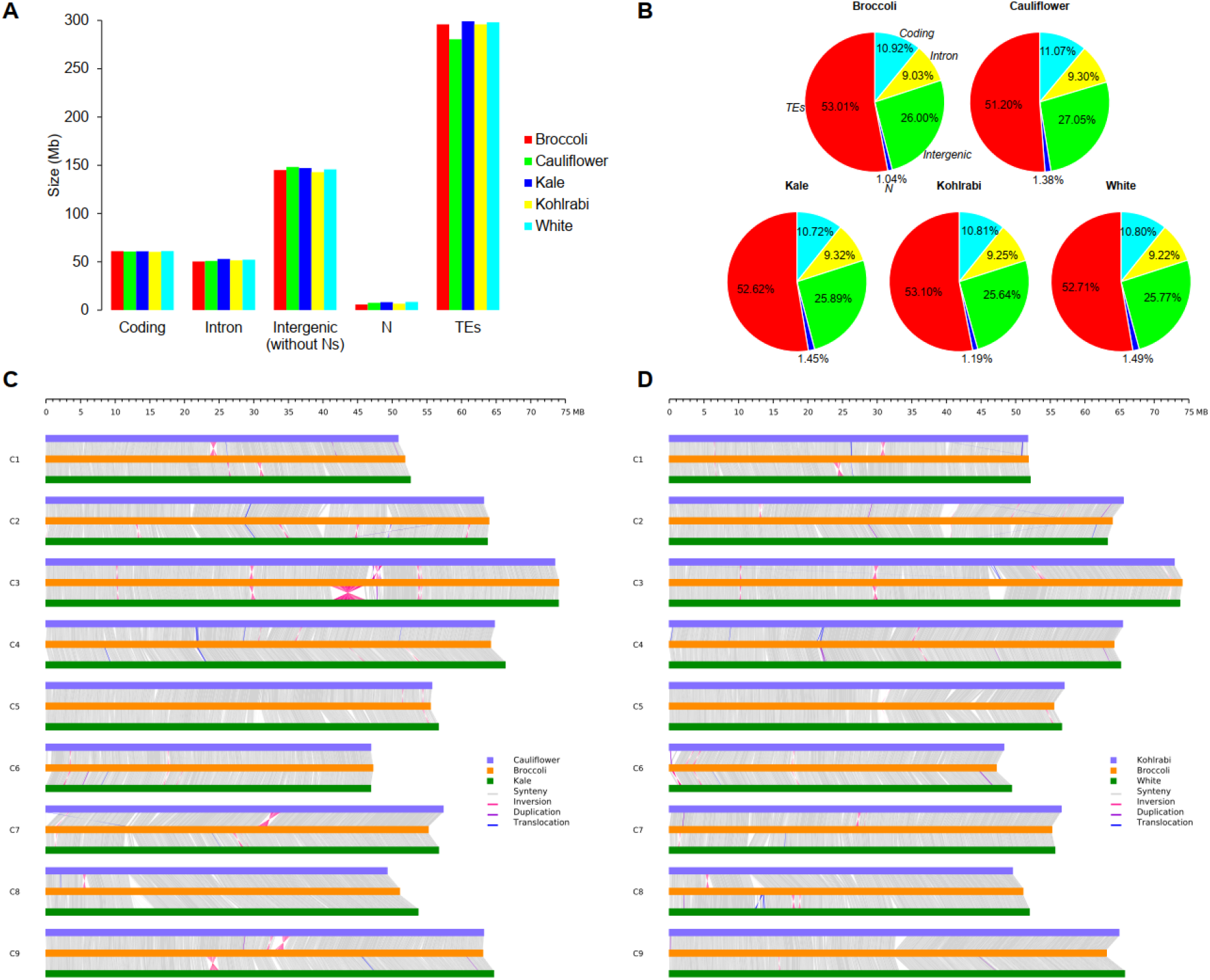
Comparisons of genomic components and whole genomic sequences among the five new *B. oleracea* assemblies. (**A**) Size of genomic components in the five assemblies. (**B**) Percentage of genomic components in the five assemblies. As indicated in (**B**) for the broccoli genome, different colours in each of the other four genomes represent different genomic components (red: TEs, aqua: coding sequence, yellow: intron sequence, green: intergenic sequences, blue: Ns). (**C**) Whole-genome comparison among cauliflower, broccoli and kale. (**D**) Whole-genome comparison among kohlrabi, broccoli and white cabbage. Note: in (**C**) and (**D**), only genomic rearrangements with length > 50Kb are shown.

On a genome-wide scale, protein-coding genes were enriched in chromosome arms while TEs tended to be enriched in (peri-)centromeric regions (Fig. S1). Overall, the five assemblies showed similar genomic components, with approximately 11% (60.25-61.09 Mb) coding sequences, 9% (50.42-52.14 Mb) intron sequences, 26% (142.89-148.19 Mb) intergenic sequences and 1% (5.80-8.44 Mb) gaps (N’s). However, the size of TEs were more variable among the five assemblies. In cauliflower genome, 280.53 Mb sequences were annotated as TEs, which was 15.33-18.52 Mb less than the figures of the other four genomes (Fig. 1A and 1B).

Comparative analyses among the five genomes show that they are highly collinear, which was demonstrated by their highly conserved genomic sequences, as well as gene content and order along each chromosome (Fig. 1C and 1D, Fig. 2, Fig. S2 and S3). Approximately 80% of each of the other four genome sequences matched in one-to-one syntenic blocks with the broccoli genome (Table S32). Substantial collinearity notwithstanding, we did detect 6.52-13.96 Mb inversions, 25.79-34.88 Mb translocations, 7.97-14.22 Mb insertions and 20.80-32.33 deletions (size ⩾ 30bp) in whole genome comparisons of each of the other four genomes with the broccoli genome (Fig. 1C and 1D, Table S33-S44). Notably, we identified eight large inversions with sizes ranging from 134 Kb to 4.88 Mb, which were verified by Bionano maps (Table S24, Fig. S16). Macrosynteny analysis also revealed five large inversions (size: 0.63 to 4.75 Mb) that were already identified based on whole genome comparison among the five genomes (Fig. 2C). Each pair of the five genomes shows clear 3:3 synteny patterns, with each region in one genome having up to three syntenic regions in another genome, indicating that *B. oleracea* genomes share a whole genome triplication (WGT) event (Fig. 2A and 2B, Fig. S2, Fig. S3). To detect genes that might be under selection, we calculated the rate of synonymous mutations per synonymous locus (Ks) between 1:1 orthologous genes for each pair of the five genomes. Comparable Ks distributions were identified between the ten comparisons with only one peak being found, suggesting that these genes may have diverged from the common ancestors of *B. oleracea* (Ks: 0.008~0.0111) (Fig. 2D). We then calculated the Ka/Ks ratios to find genes with evidence for selection. Because most nonsynonymous mutations are deleterious and experienced strong purifying selection (Sun et al., 2018), the Ka/Ks ratios for most orthologous genes are expectedly close to zero. Approximately 13,726~17,003 genes in each of the five genomes were identified likely to be under strong purifying selection (Ka/Ks<0.6). In contrast, relatively few genes (1,737-2,038) were detected to be under positive selection (Ka/Ks>1) (Fig. 2E).

**Fig. 2.**
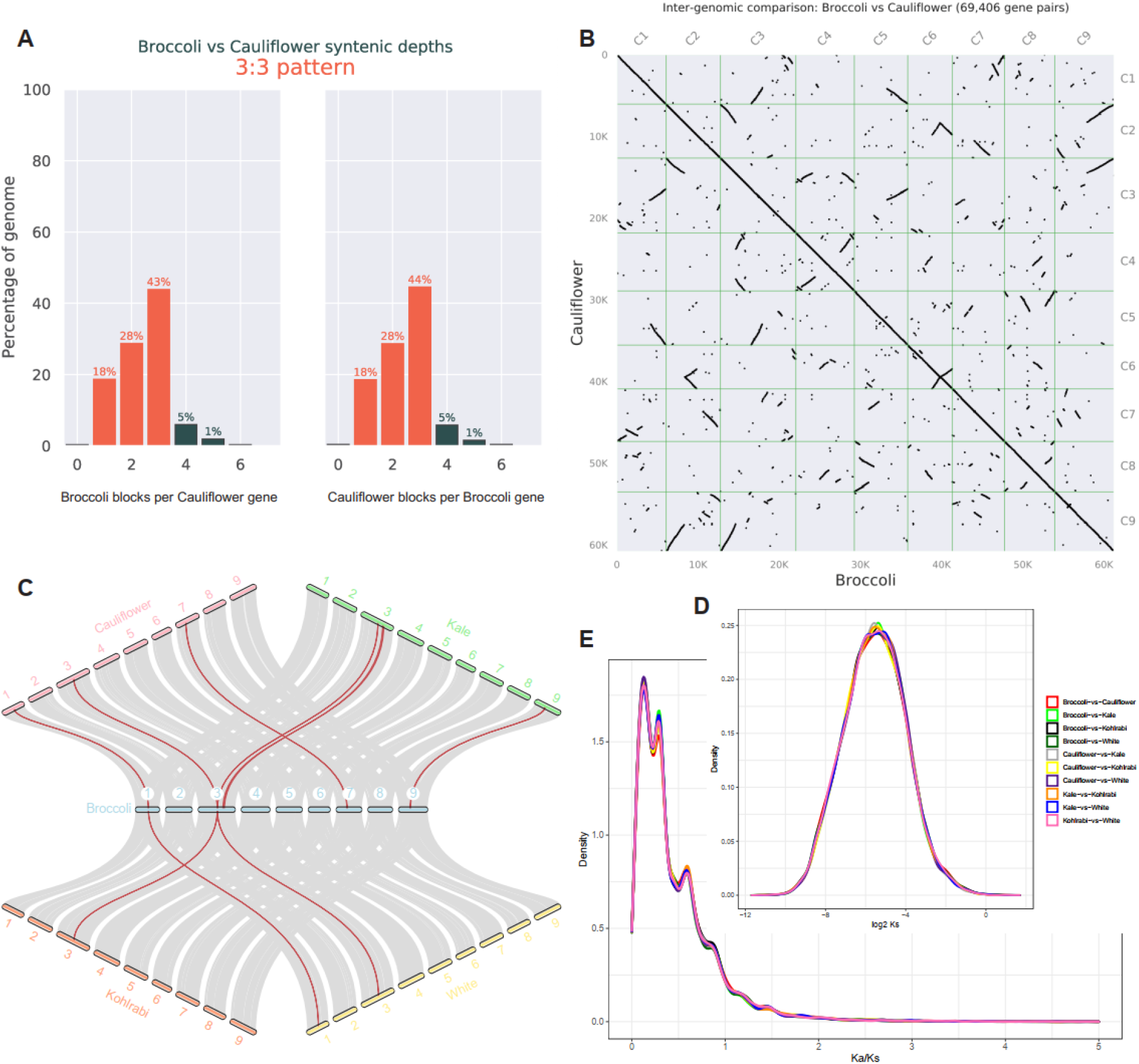
(**A**) Ratio of syntenic depth between broccoli and cauliflower genome. (**B**) Homologous dot plot between broccoli and cauliflower genome. (**C**) Macrosynteny among the five genomes with grey links connecting blocks of >30 one-to-one gene pairs. Red lines indicate inversions. (**D**) Ks distribution of each two of the five genomes. (**E**) Ka/Ks distribution of each pair of the five genomes.

### Composition and features of *B. oleracea* pan-genome

We constructed a *B. oleracea* pan-genome comprising our five new assemblies and four published genomes (Belser et al., 2018; Cai et al., 2020; Guo et al., 2021), all of which were assembled using long-read sequencing technology. Pairwise whole genome and gene set comparisons, and modelling of the pan-genome suggested a pan-genome size of ~653 Mb with ~84,000 genes and a core-genome size of ~458 Mb including ~33,000 genes. Both pan-genome size and pan-gene number increased when adding additional genomes, however both the core-genome size and core-gene number decreased (Fig. 3A and 3B). Using OrthoFinder, we detected 60,940 gene clusters in the *B. oleracea* pan-genome, consisting of 535,182 annotated genes in the nine genomes. Of these total gene clusters, 37,669 (~61.81%), 21,891 (~35.92%) and 1,380 (~2.27%) were considered as core, dispensable and specific gene clusters, respectively (Fig. 3C and 3D). The proportion of core gene clusters was much higher than the percentages for dispensable and specific gene clusters. We found 48,543-49,303 core genes representing 79.53%-80.05% of total gene models in our five newly generated genomes. A total of 62-117 specific genes and 530-707 orphan genes (genes with no paralogues and no orthologues in other genomes) were detected in the five genomes.

**Fig. 3.**
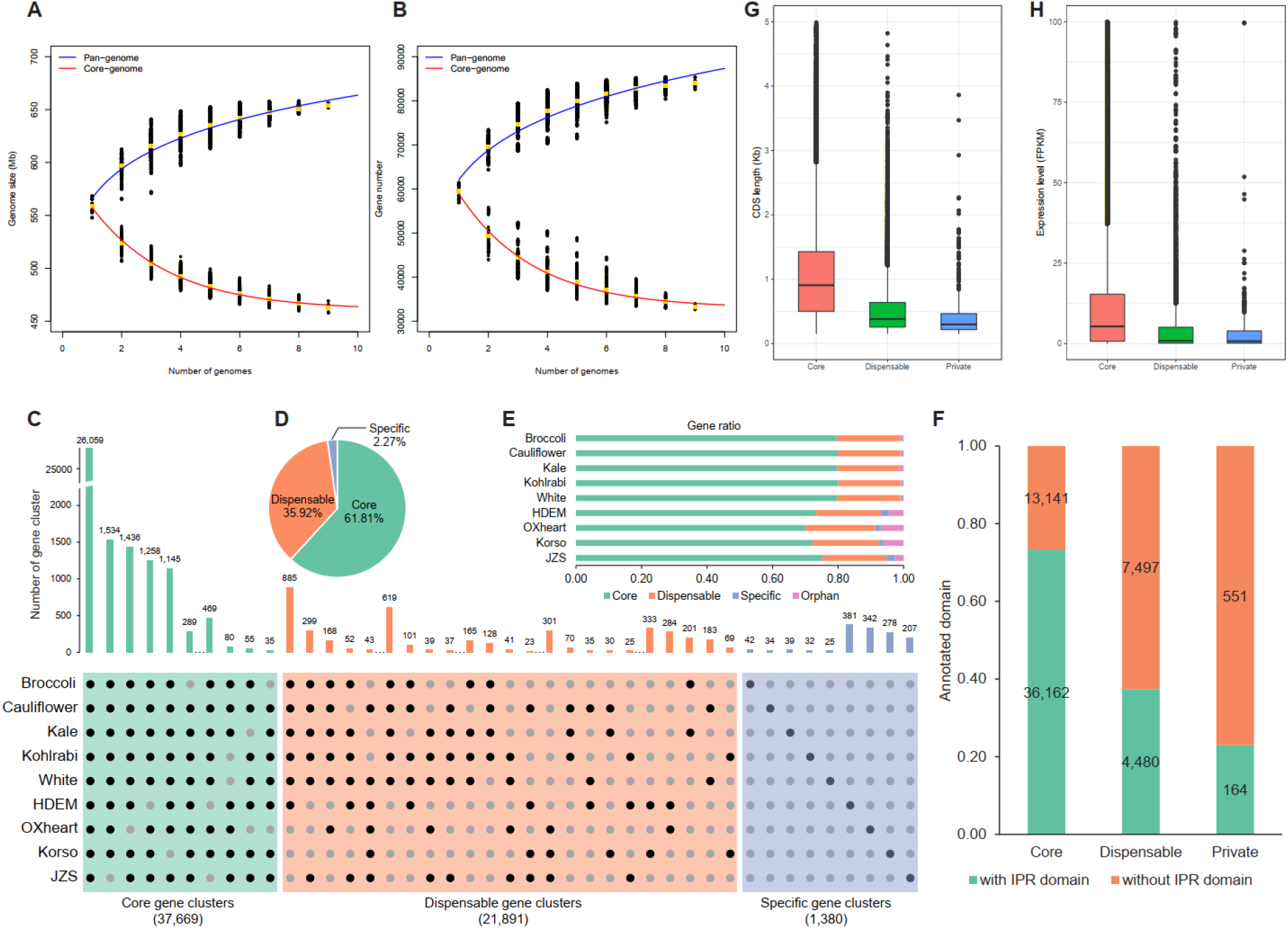
Pan-genome analyses of nine *B. oleracea* genomes. (**A**) Pan-genome and core-genome modelling for sequences, which were based on pairwise whole-genome comparisons across nine genomes. (**B**) Pan-genome and core-genome modelling for gene space, which was based on gene set comparisons across nine genomes. (**C**) Compositions of the *B. oleracea* pan-genome. The histograms show subsets of core, dispensable and specific gene clusters. The dots in between the histogram bars refer to other combinations that are not displayed (see Table S45). (**D**) Ratio of gene clusters marked by each composition. (**E**) Ratio of classified genes in each genome. Orphan genes are those with no paralogues and no orthologues in other genomes. (**F**) Proportion of genes with InterPro (IPR) domains in the Broccoli core genes, dispensable genes and private genes. Private genes include genes within specific gene clusters and orphan genes. (**G**) CDS length of core, dispensable and private genes in Broccoli genome. (**H**) Expression level of core, dispensable and private genes in Broccoli genome.

We found that 73.35% of the core genes in the broccoli genome were annotated with InterPro domains. This proportion was much higher than the percentages for the dispensable and private genes (including specific genes and orphan genes), which accounted to 37.41% and 22.94% resp. (Fig. 3F). The average CDS length of core genes are significantly longer than those of less conserved genes (Student-Newman-Keuls test with *a* = 0.01) (Fig. 3G). Additionally, the average gene expression level of the core genes was significantly higher than those of dispensable and private genes (Student-Newman-Keuls test with *a* = 0.01) (Fig. 3H). Gene Ontology (GO) enrichment analysis showed that core genes were mainly enriched in essential biological processes, including regulation of metabolic processes, biosynthetic process and transcription, host cellular component and nucleotide binding process (Fig. S4A). However, dispensable genes were mainly enriched in terms of cellular respiration, translation, organelle component and ribonuclease activity. Kyoto Encyclopedia of Genes and Genomes (KEGG) pathway analyses showed that core genes were enriched in pathways related to transcription factors, specific metabolism, transporters, replication and repair, whereas dispensable genes were mainly enriched in pathways related to ribosome, translation, photosynthesis, oxidative phosphorylation (Fig. S4B).

### Different evolutionary dynamics of LTR-RTs in *B. oleracea* and *B. rapa*

We detected 4,739-5,995 intact LTR-RTs, totalling 32.04-40.57 Mb in each of the nine *B. oleracea* long-read assemblies, however, only 690 and 1,482 (3.42 and 8.13 Mb) intact LTR-RTs were found in two short-read assemblies (JZSv1 and TO1000) (Table S19). The remarkably less intact LTR-RTs in short-read assemblies could be due to the collapse of short reads from these regions. Copia-like LTR-RTs are more abundant than Gypsy-like LTR-RTs in all *B. oleracea* genomes (Fig. 4A). The nine long-read assemblies showed similar LTR-RTs length distribution. Generally, Copia-like LTR-RTs peaked at around 5-6Kb, while the lengths of Gypsy-like LTR-RTs were more variable (Fig. S5). On average, the lengths of Gypsy-like LTR-RTs are longer than those of Copia-like LTR-RTs. In the published 18 *B. rapa* genomes, we only identified 2,573-3,958 intact LTR-RTs, totalling 17.19-26.33 Mb (Fig. 4A and 4B, Table S20). Statistically, *B. oleracea* accumulated significantly more LTR-RTs than *B. rapa* (Fig. 4A and 4B), consistent to a previous study (Liu et al., 2014a). The length distributions of LTR-RTs among the 18 *B. rapa* genomes are similar (Fig. S6), which are also comparable to those of *B. oleracea*.

**Fig. 4.**
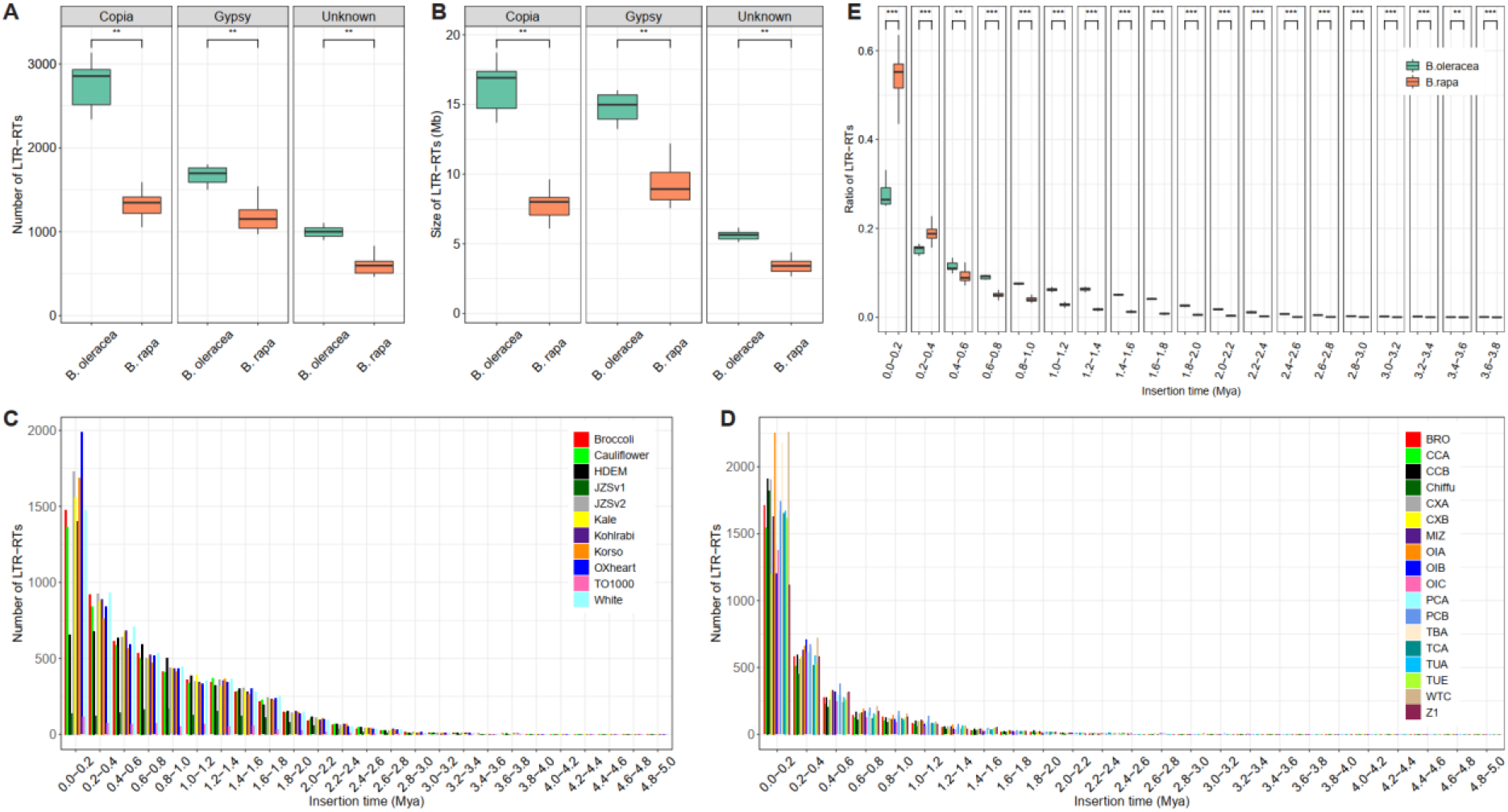
Evolutionary dynamics of full-length LTR-RTs in *B. oleracea* and *B. rapa*. The number (**A**) and total genome size (**B**) of full-length LTR-RTs in *B. oleracea* and *B. rapa* pan-genome. Distributions of insertion time dated by divergence rate of full-length LTR-RTs (the rate of neutral mutation sites accumulated in the two terminal repeat sequences of LTR-RTs). This was calculated in the (**C**) 11 genome assemblies of *B. oleracea* and (**D**) 18 genome assemblies of *B. rapa*. (**E**) Ratio of LTR-RTs that were accumulated in each time range in *B. oleracea* and *B. rapa* genomes. JZSv1 and TO1000 were not included in this calculation. Significance is determined by a two-sided Wilcoxon rank-sum test.

Analysis of LTR insertion time revealed continuous LTR-RTs expansion in all nine long-reads assembled *B. oleracea* genomes since ~3 Mya (Fig. 4C), after the speciation with its sister species of *B. rapa* (Cheng et al., 2017). We identified two LTR-RTs burst events in *B. oleracea* genomes, a “young” event that has taken place in eight long-read assembled genomes (not present in HDEM) around 0-0.2 Mya, and an “old” event that occurred in all nine genomes around 1.2-1.4 Mya, with 25.07%-33.14% and 5.58%-6.90% LTR-RTs being formed at these two time ranges, respectively (Fig. 4E). Generally, the evolutionary dynamics of LTR-RTs in *B. oleracea* genomes are similar. Nevertheless, the remarkable LTR-RTs number variation, especially for the newly inserted LTR-RTs, among *B. oleracea* genomes may indicate different changes in LTR-RTs patterns during the intraspecific diversification. As an illustration, we found less than twice as many newly inserted LTRs in HDEM than in the other eight genomes. In *B. rapa*, continuous LTR-RTs expansion initiated later, at ~2 Mya (Fig. 4D). Only one LTR-RTs burst event was found in each of the 18 *B. rapa* genomes, which is shared in time with the “young” LTR-RTs burst event in *B. oleracea*, with significantly higher percentage of LTR-RTs (43.53%-63.55%) being accumulated during 0-0.2 Mya than in *B. oleracea* (Fig. 4E). In *B. rapa*,the vast majority of LTR-RTs (66.30%-79.22%) were formed at the time range of 0-0.4 Mya, whereas only 28.23%-47.16% LTRs were accumulated in *B. oleracea* at the same time range (Fig. 4E). *B. oleracea* accumulated significantly higher percentage of LTRs than *B. rapa* from ~3.8 Mya until ~0.4 Mya (Fig. 4E). Together, these data suggested that LTRs accumulated faster in the recent ~0.6 Mya and later in *B. rapa* than in *B. oleracea*, indicating different evolutionary dynamics of LTR-RTs in the two sister species.

### Faster gene loss in *B. rapa* than in *B. oleracea* before intraspecific diversification

Using a pan-genome strategy (Cai et al., 2021), we inferred the ancestral genome of *B. oleracea* (A_Bol_). A total of 33,287 WGT-derived genes (14,153, 10,192, and 89,42 genes in LF, MF1 and MF2 subgenomes, respectively) formed A_Bol_, 3,121 more than in the inferred *B. rapa* ancestral genome (A_Bra_) (Cai et al., 2021). Sliding windows with 500 genes and an increment of two genes were used to calculate gene densities in the three subgenomes. Similar to its sister genome A_Bra_, we discovered the phenomenon of subgenome dominance in ABol, with average gene densities of 0.745, 0.537 and 0.470 in LF, MF1 and MF2 subgenomes respectively (Fig. S7A). Additionally, using the broccoli genome as a representative of the diverse morphotypes, we found significantly lower gene densities in all its three subgenomes compared to ABol. On average, 0.075, 0.091 and 0.110 genes in broccoli LF, MF1 and MF2 subgenomes were fractionated (Fig. S7B). The gene density and gene fractionation distributions along the seven AKBr chromosomes among the nine *B. oleracea* extant genomes were similar (Fig. S12 and S13), consistent with the broccoli representative genome. Our results suggest that genes were extensively fractionated in all the individual genomes of *B. oleracea* during the intraspecific diversification (Fig. S13). Gene fractionation patterns along 24 ancestral karyotype (AK) blocks for nine *B. oleracea* genomes representing diverse morphotypes were also comparable in all three subgenomes, with blocks D and G of the LF subgenome showing much higher levels of gene loss (Fig. S8). Most *B. rapa* genomes, also representing diverse morphotypes, also showed similar gene fractionation patterns along the seven AKBr chromosomes as well as the 24 AK blocks (Fig. S9, Fig. S14, Fig. S15), which were also generally in agreement with those of *B. oleracea*. However, gene fractionation ratios in specific blocks for several *B. rapa* genomes differed remarkably from the other *B. rapa* genomes, such as blocks G, N, T and V of OIC’s (*ssp. oleifera*, Rapid cycling’s) LF subgenome, block I of TUE’s (*ssp. rapa*, European Turnip’s) MF1 subgenome, block V of CXA’s (*ssp. parachinensis*, Caixin’s) MF2 subgenome, and etc (Fig. S9). Interestingly, the blocks D and G showed increased gene fractionation in the LF subgenome of almost all *B. rapa* and *B. oleracea* morphotypes.

We further constructed the common ancestral genome of *B. oleracea* and *B. rapa* (A_Bol_Bra_), a gene repertoire consisting of 34,547 WGT-derived genes (14,539, 10,659 and 9,349 in LF, MF1 and MF2, respectively), by merging genes in A_Bol_ and A_Bra_ and ordering the non-redundant genes in the tPCK karyotype. The average gene densities relative to A_Bol_Bra_ for all three subgenomes of A_Bol_ (0.745, 0.537 and 0.470 in LF, MF1 and MF2, respectively) were significantly higher than those of A_Bra_ (0.727, 0.507 and 0.435 in LF, MF1 and MF2, respectively) (Fig. 5A). In agreement with this, the average gene fractionation ratios in three subgenomes of A_Bol_ (0.027, 0.044 and 0.045 in LF, MF1 and MF2, respectively) were significantly lower than those of A_Bra_ (0.101, 0.141 and 0.165 in LF, MF1 and MF2, respectively) (Fig. 5B), which was also reflected in the 24 AK blocks (Fig. S10). This suggests that after the speciation between *B. oleracea* and *B. rapa*, WGT-derived gene loss in *B. rapa* was stronger than that in *B. oleracea*. We further investigated gene fractionation in individual genomes of *B. oleracea* and *B. rapa*. In contrast to the different fractionation rates of their respective ancestral genomes, similar average gene fractionation ratios were identified between the 18 genomes of *B. rapa* (0.084-0.129 in LF, 0.108-0.174 in MF1 and 0.112-0.165 in MF2, respectively) and the nine genomes of *B. oleracea* (0.075-0.105 in LF, 0.091-0.124 in MF1 and 0.104-0.128 in MF2, respectively) (Fig. 6A and 6B).

**Fig. 5.**
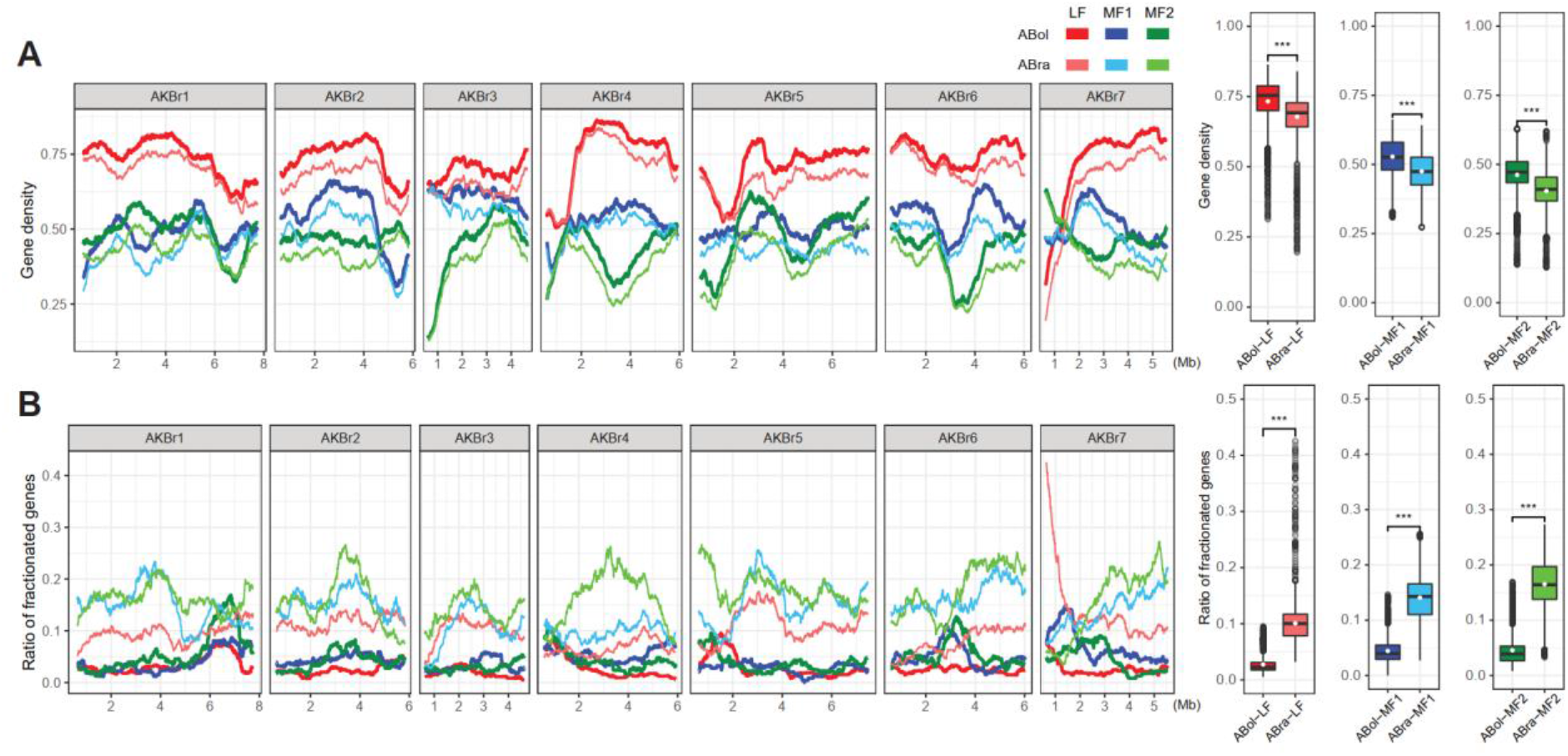
Less gene fractionation in ancestral genome of *B. oleracea* (A_Bol_) than in ancestral genome of *B. rapa* (A_Bra_). (**A**) Gene density distribution on the seven inferred chromosomes of AKBr in the three subgenomes of A_Bol_ and A_Bra_. The figure on the right shows gene density in each window. Two-tailed Student’s t-test was performed to compare gene densities between A_Bol_ and A_Bra_ for each subgenome. (**B**) Gene fractionation distribution in the three subgenomes of A_Bol_ and A_Bra_. The figure on the right shows gene fractionation ratio in each window. Two-tailed Student’s t-test was performed to compare gene fractionation ratio between A_Bol_ and A_Bra_ for each subgenome. 500-gene windows with an increment of two genes was used to calculate gene density and gene fractionation ratio in (**A**) and (**B**).

**Fig. 6.**
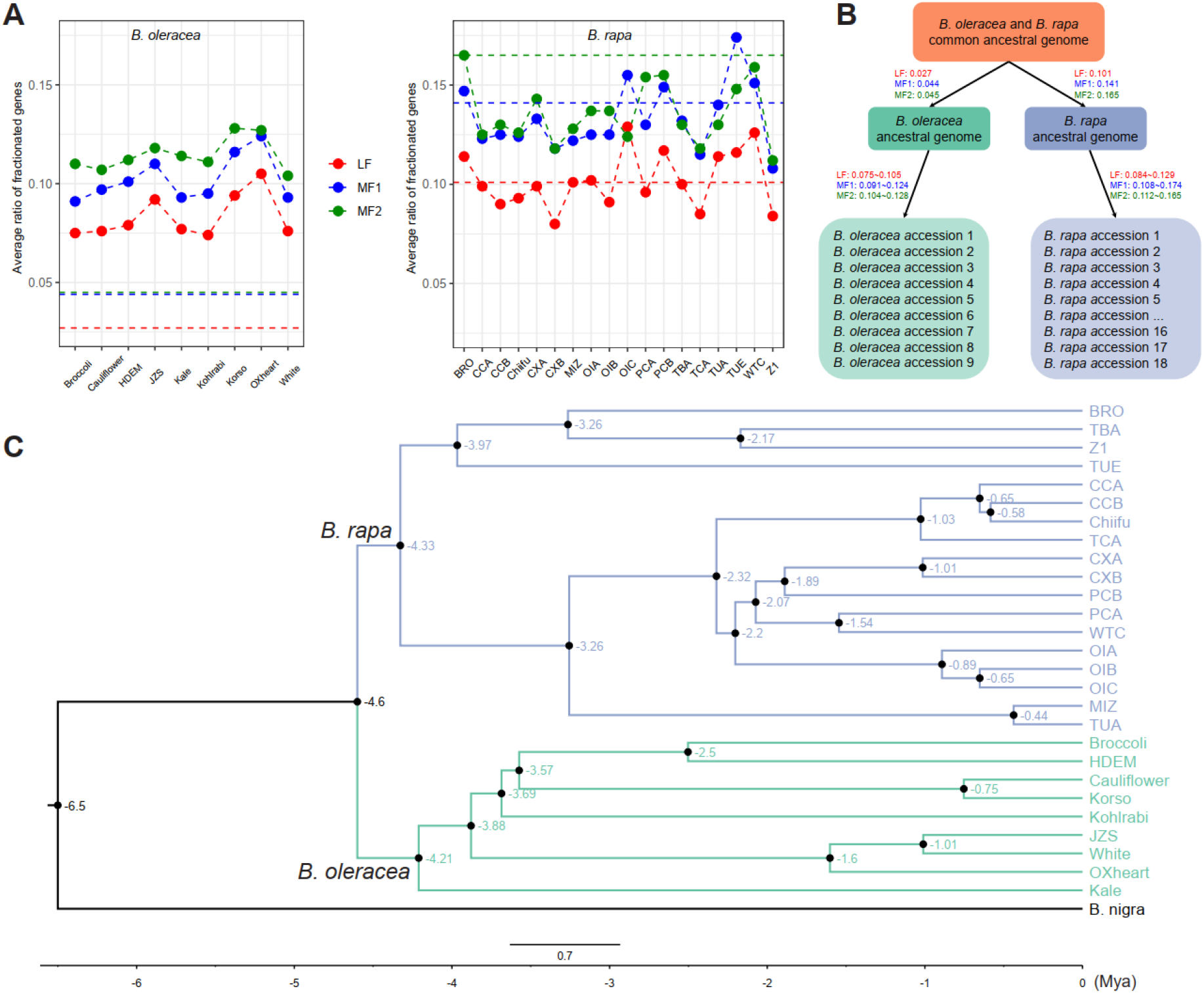
Different gene fractionation patterns between *B. oleracea* and *B. rapa*. (**A**) Average gene fractionation ratio of nine *B. oleracea* and 18 *B. rapa* accessions in the three subgenomes. Red, blue and green horizontal dashed lines represent average gene fractionation ratios of ancestral genome of *B. oleracea* (left) and ancestral genome of *B. rapa* (right) relative to A_Bol_Bra_. (**B**) Gene fractionation patterns from A_Bol_Bra_ to extant genomes of *B. oleracea* and *B. rapa*. Values next to arrows represent average gene fractionation ratios from the ancestral genome to its descendant genome in the three subgenomes. (**C**) Phylogenetic tree of 28 *Brassica* genomes and their estimated divergence times (million years ago).

A phylogenetic tree including the genomes of 18 *B. rapa*, nine *B. oleracea* and one *B. nigra*(outgroup) was constructed using 4,756 single-copy orthologous genes. We revealed two main lineages: one for *B. oleracea* and the other one for *B. rapa*, both of which were supported by high bootstrap values (Fig. S11). Within the *B. oleracea* lineage, we revealed the two main cultivated lineages: “Arrested Inflorescence Lineage (AIL)” and “Leafy Head Lineage (LHL)”, with kohlrabi situated at the junction of these two lineages, consistent with an earlier report (Cai et al., 2022a). Our phylogenetic tree also supported the hypothesis that kale lineage leads to the “AIL” and “LHL” (Cai et al., 2022a) (Fig. 6C, Fig. S11). The genealogical relationships within *B. oleracea* were consistent with morphotypes rather than geographical origin. The *B. rapa* lineage in our analysis largely corroborated the one revealed by Cai et al (Cai et al., 2022b) with minor differences, both of which revealed five monophyletic groups for these diverse *B. rapa* morphotypes. Estimation of divergence times based on Bayesian inference using MCMCTree program suggested that intraspecific diversification for the sister species *B. oleracea* (~4.21 Mya) and *B. rapa* (~4.33 Mya) happened simultaneously and shortly after their speciation (~6.5 Mya) (Fig. 6C). Together with the gene fractionation ratios as observed in this study, these results suggest that *B. rapa* experienced faster WGT-derived gene loss than *B. oleracea* before their intraspecific diversification, however, during the intraspecific diversification, *B. oleracea* and *B. rapa* experienced gene loss at a comparable speed.

### Biased gene loss within and between *B. oleracea* and *B. rapa*

A total of 1,260 (386, 467 and 407 in LF, MF1 and MF2, respectively) WGT-derived genes were lost in the three subgenomes of A_Bol_ in relation to A_Bol_Bra_ (Fig. 7A). Among these non-redundant genes, we observed that 97.2% lost one copy of their paralogues, 2.6% lost two copies and 0.2% three copies, suggesting a strong bias towards losing only one copy among the three paralogous genes. In comparison, much more genes were lost in A_Bra_, with 1,423 1,477 and 1,481 genes in LF, MF1 and MF2, respectively. A total of 71.1% non-redundant genes lost only one copy of the three paralogous genes in A_Bra_, again suggesting biased loss of only one paralogous copy. Interestingly in A_Bra_, we observed a surprisingly high ratio (26.1%) of non-redundant genes that had lost all the three copies of paralogous genes (Fig. 7B), which is a major factor contributing to the faster gene loss in A_Bra_ than A_Bol_. Gene ontology (GO) enrichment analysis of these genes of which all three copies were lost (738 genes) showed that the vast majority of enriched GO terms were associated with biological process, including biological regulation, response to stimulus, response to chemical, response to phytohormones (auxin, cytokinin and jasmonic acid) and etc (Table S25), which suggests a large amount of ancestral *B. rapa*-specific relaxations of biological or environmental constraints leading to co-elimination of all three copies of paralogous gene. In addition, the two species also showed biased patterns of gene loss with respect to the lost gene functions. Enriched GO terms of lost genes in A_Bol_ were mainly related with ‘housekeeping roles’, such as DNA damage response, ATP/ADP binding, nuclease activity and other essential cellular functions (Table S26-S28). In A_Bra_, besides functional GO categories with ‘housekeeping roles’, the vast majority of enriched GO terms of the lost genes belong to biological process including those likely related to the adaptations to changes in environmental conditions (Table S29-S31). This suggests that A_Bra_ suffered relaxation of biological or environmental constraints more so than A_Bol_, indicating species-specific adaptation to the environment through adaptive gene loss. During intraspecific diversification, the extant genomes of *B. oleracea* and *B. rapa* both showed clear bias to one-copy gene loss and remarkably few three-copy gene loss (Fig. 7C and Fig. S17). We then investigated gene loss bias between each two extant individual genomes of *B. oleracea* or *B. rapa*. Among all *B. oleracea* pairwise genome combinations, we found that on average only 33.42%, 37.43% and 37.81% common genes were lost in LF, MF1 and MF2, respectively (Fig. 7D, Fig. S18). These figures were relatively higher in *B. rapa*, with 38.88%, 43.38% and 44.88% in LF, MF1 and MF2, respectively (Fig. 7D, Fig. S19). These results suggest that gene loss remains biased among extant genomes during the intraspecific diversification. Among all pairwise genome comparisons between extant *B. oleracea* and *B. rapa* genomes, we found on average only 11.25%, 11.49% and 11.89% shared gene loss in LF, MF1 and MF2, respectively (Fig. 7E, Fig. S20). Taken together, our results suggest the continuing gene loss bias, both within and between species, during intraspecific diversification of *B. oleracea* and *B. rapa*.

**Fig. 7.**
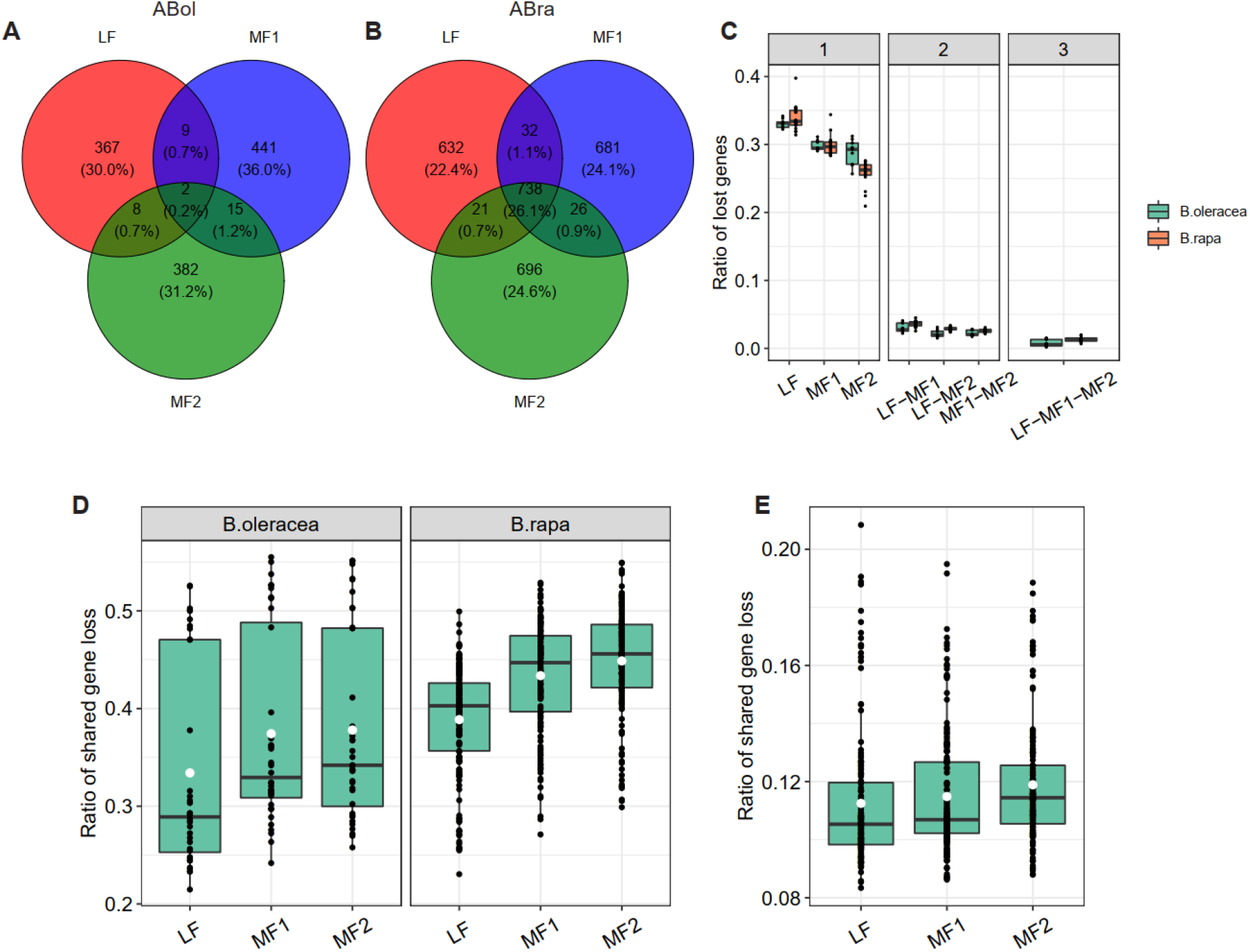
Biased gene loss in *B. oleracea* and *B. rapa*. Common and unique gene losses among the three subgenomes of A_Bol_ (**A**) and A_Bra_ (**B**). (**C**) The ratio of one-copy, two-copy and three-copy gene loss in the extant genomes of *B. oleracea* or *B. rapa*. Each black dot represents one individual genome. (**D**) Distribution of shared gene loss between each two of the extant genomes of *B. oleracea* or *B. rapa* in the three subgenomes. Each black dot represents one pairwise combination of the nine *B. oleracea* or 18 *B. rapa* genomes. (**E**) Distribution of shared gene loss between extant genomes of *B. oleracea* and *B. rapa* in the three subgenomes. Each black dot represents one pairwise combination of the nine *B. oleracea* and 18 *B. rapa* genomes. White dots indicate the average value in (**D**) and (**E**).

## Discussion

In this study, we *de novo* assembled chromosome-scale reference genomes for five different *B. oleracea* morphotypes by integrating data from short-read sequencing (Illumina), long-read sequencing (Oxford Nanopore and Pacific Biosciences) and Bionano Genomics DLS optical maps. To our knowledge, these five assemblies exhibit the highest contiguity among released *B. oleracea* genomes. Comparative analysis revealed both highly syntenic relationships and extensive structural variants among the five genomes, which highlights the insufficiency of single-reference genomes to represent the sequences of a species. These five newly generated assemblies together with other published high-quality *B. oleracea* reference genomes provide an opportunity to investigate the composition and features of *B. oleracea* pan-genome via a *de novo* assembly approach (Danilevicz et al., 2020). Moreover, *B. oleracea* is one of the ideal models for studying polyploidization and evolution in plants. With these genomes representing diverse morphotypes, we inferred the ancestral genome of *B. oleracea* using a pan-genome strategy (Cai et al., 2021) and systematically studied the WGT-derived gene fractionation during its intraspecific diversification, as well as compared the patterns with its sister species *B. rapa*. The present work not only provides valuable genomic resources to the Brassica scientific community for *B. oleracea* improvement, but also provides insights towards understanding individual genome evolution during the intraspecific diversification of *B. oleracea* and *B. rapa*.

The completeness of a reference genome is of great importance to reliably detect LTR elements with complex structures. In the present study, we used the same approach to identify intact LTR-RTs in 11 *B. oleracea* genomes (nine long-read and two short-read assemblies, respectively) and 18 *B. rapa* genomes. By comparing the LTR-RTs content between the nine long-read and the two short-read *B. oleracea* assemblies, we showed that much more intact LTR-RTs were detected in long-read assemblies, with the sizes being 9.37-11.86 times larger than TO1000 genome and 3.94-4.99 times larger than JZS v1 genome. More complete detection of LTR-RTs also provided new insights into the evolution of LTR-RTs in *B. oleracea*. A “young” LTR outbreak event was identified in eight *B. oleracea* genomes, however this event was not observed in the two short-reads assemblies probably because these LTR-RTs were not successfully assembled in these two genomes. This “young” LTR outbreak is also not identified in the HDEM genome, however, we cannot rule out whether this is due to assembly artifacts. Transposable elements (TEs) have played an important role in the evolution of plant genomes (Cai et al., 2020; Chuong et al., 2017; Lisch, 2013; Liu et al., 2021a). Among different categories of TEs, LTR-RTs have been shown to contribute significantly to genome size expansion in plants owing to their high copy number and large size (Ming et al., 2015; Nystedt et al., 2013; Ou and Jiang, 2018; Rensing et al., 2008; Schnable et al., 2009). It is reported that LTR-RTs are highly unstable in plant genomes (Devos et al., 2002; Domansky et al., 2000; Liu et al., 2021a; SanMiguel et al., 1998), and it is often observed that LTR-RT components vary among different subspecies (Cai et al., 2020; Sun et al., 2022). Our results also showed that LTR-RTs components strongly differed in both *B. oleracea* and *B. rapa* genomes, suggesting LTR-RTs as important drivers for intraspecific diversification. Assuming that *B. oleracea* and *B. rapa* diversified at ~4.6 Mya (Cheng et al., 2017), we would conclude that nearly all LTR-RTs in these two species were inserted during the period of intraspecific diversification (Fig. 4A and 4B) and no LTR-RTs were inherited from their common ancestor. A burst of transposable elements has been reported in connection with taxonomic groups and species formation as well as domestication (Belyayev, 2014). We see different patterns of both LTR-RTs and gene loss dynamics in *B. rapa* and *B. oleracea* extant genomes, which should be studied in depth to understand the diversification history of these two species.

Single-reference genomes are not sufficient to cover the whole genome sequence of a species. This has been confirmed by pan-genome studies in major crops, such as soybean (Liu et al., 2020), rice (Qin et al., 2021), maize (Hufford et al., 2021) and rapeseed (Song et al., 2020). Previously, a *B. oleracea* pan-genome including nine varieties and a wild relative was constructed using an iterative assembly approach with NGS data. Even though nearly 18.7% of the pan-genome is composed of variable genes, it is still likely that genomic variations resolved in this pan-genome are underestimated. First reason is that numerous complex variations cannot be detected by simply mapping short reads to the reference genome (Liu et al., 2020). Second reason is that short reads often result in incomplete assemblies. Indeed, in our research, we found ~35.92% dispensable gene clusters and ~2.27% specific gene clusters in the pan-genome constructed based on high-quality *de novo* assembled sequences, which have the potential to resolve the vast majority of genomic variations. More than 20% of all genes in each individual genome were assigned as dispensable, specific or orphan, suggesting that SVs widely exist between different morphotypes within *B. oleracea*. Pan-genome modelling suggested a *B. oleracea* pan-genome size of ~653 Mb and a core-genome size of ~458 Mb. *B. oleracea* includes many morphotypes with enormous phenotypic diversity, however in this pan-genome analysis the nine genomes cover only five morphotypes. Inclusion of more morphotypes or increasing the number of samples likely leads to a larger estimate of the pan-genome size. Our five new assemblies are nearly identical in terms of ratio between each of the core, dispensable, specific and orphan genes (Fig. 3E). However, these figures strongly differ among the four published genomes which were generated by different labs with different approaches (Fig. 3E). This inconsistency might have been induced by different genome assembly and gene prediction approaches. To minimize such effects in a pan-genome study, by best solution is to include a large panel of samples in a single project to capture the whole pan-genome and implement same approach for assembly and gene model prediction, which is however costly. Nevertheless, the pan-genome dataset constructed from this study is valuable for important functional genes and genetic breeding studies as it provides comprehensive genomic variations in the gene pool for *B. oleracea*.

*B. oleracea* and *B. rapa* are both mesopolyploid species that have been domesticated into a remarkable variation in morphotypes, with genomes that have experienced a triplication event, followed by extensive gene fractionation and chromosome rearrangements (Cheng et al., 2012a). In *B. rapa*, gene fractionation during individual genome evolution has been investigated by constructing an inferred ancestral genome based on a pan-genome approach. Cai and colleagues observed the continuing influence of the dominant subgenome on *B. rapa* intraspecific diversification (Cai et al., 2021). The *B. oleracea* pan-genome present in this study allowed us to infer the ancestral genome of *B. oleracea*, as well as a common ancestral genome of *B. oleracea* and *B. rapa*. Instead of using a single reference genome for each species, we constructed the common ancestral genome by including nine *B. oleracea* and 18 *B. rapa* genomes. This remarkably improved the common ancestral genome since gene content is added from diverse *B. oleracea* and *B. rapa* genomes, as was discussed by Cai et al (Cai et al., 2021). The three inferred ancestral genomes provide the opportunity to systematically study gene fractionation patterns both before and during intraspecific diversification of the two species. Our data suggest that the ancestral genome of *B. rapa* has undergone stronger gene fractionation than the ancestral genome of *B. oleracea*. Based on divergence time estimation, the ancestors of *B. oleracea* and *B. rapa* had similar subspeciation initiation times. This brings us to conclude that the ancestral genome of *B. rapa* has experienced faster gene loss than *B. oleracea*. This largely attributes to the extensive loss of all three -copies of many genes in the *B. rapa* ancestral genome. We hypothesize that the ancestral *B. rapa* has undergone its specific relaxations of given biological or environmental constraints, which result in the ‘co-elimination’ of all the three copies of redundant genes that were triplicated by the WGT. As expected, we observed the continuing influence of the dominant subgenome on intraspecific diversification in *B. oleracea*, similar as in *B. rapa*. Different from the diversification of the two ancestral genomes, extant genomes of the two species show comparable speed of gene loss during their intraspecific diversification, which is likely driven by human domestication. Our data support the finding that gene loss is biased towards both genomic position and function. Indeed, the dominant LF subgenome displays significantly more genes than MF1 and MF2 subgenomes. The lost genes of ancestral genomes of *B. oleracea* and *B. rapa* differ in functional GO categories. Furthermore, both in *B. oleracea* and *B. rapa*, there is a strong bias towards one-copy gene losses, both before and during intraspecific diversification. The large fraction of unique gene losses between their extant genomes suggests the continuing gene loss bias during intraspecific diversification of the two species.

## Materials and Methods

### Plant materials and DNA sequencing

Five *B. oleracea* accessions (DH lines) representing five different morphotypes, broccoli, cauliflower, kale, kohlrabi and white cabbage, were used for sequencing and *de novo* assembly in this study.

Young leaves were collected and high molecular weight (HMW) DNA was extracted for each plant following a previously established protocol (Murray and Thompson, 1980). The SQK-LSK109 Ligation Sequencing kit (Oxford Nanopore Technologies; Oxford, UK) was used for library constructions according to manufacturer’s instructions. Long-read sequencing data were generated using the Oxford Nanopore GridION platform and a run-time of 48 hr. Broccoli and kale samples were both sequenced using three flow cells, and cauliflower, kohlrabi and white cabbage samples were each sequenced using two flow cells. PacBio SMRTbell libraries were constructed from 5 μg HMW DNA using the SMRTbell Express Template Prep Kit v1 following the manufacturer’s protocols. Sequencing was performed using diffusion loading on the “Sequel SMRT Cell 1M v2” with “Sequel Sequencing Kit 2.1” reagents. The concentration for sequencing was set at 8pM for all samples. To each SMRTcell, a Sequel “DNA Internal Control Complex 2.1” was added at a low percentage according to the protocol.

In addition, genomic DNA was extracted from these young leaves using a cetyltrimethylammonium bromide (CTAB) method (Allen et al., 2006). Illumina libraries with ~450bp and ~600bp insertion sizes were constructed at GenomeScan, the Netherlands. The NEBNext® Ultra DNA Library Prep kit for Illumina (cat# NEB #E7370S/L) was used to process the DNA samples. Fragmentation of the DNA using the Biorupor Pico (Diagenode), ligation of sequencing adapters, and PCR amplification of the resulting product were performed according to the procedure described in the NEBNext Ultra DNA Library Prep kit for Illumina Instruction Manual. The quality and yield after sample preparation was measured with the Fragment Analyzer. The resulting libraries were sequenced on Illumina Hiseq 2500 (~450bp libraries) and X10 (~600bp libraries) platforms.

### RNA-seq sequencing

To aid in genome annotation, we generated mRNA-seq data for each of the five morphotypes. For each morphotype, whole young seedling and different tissues including leaves, meristems, curds, stems, and flowers were pooled in one mRNA-seq library (Table S21). We included young seedlings that were cultivated under normal condition and under heat treatment (35°C). We also included leaves that were under different treatments, including normal condition, heat treatment (35°C) for 7 days, drought treatment (no water) for 7 days and cold treatment (10°C) for 7 days. Five mRNA-seq libraries were sequenced by the Illumina NovaSeq platform with 150bp paired-end reads. Raw reads were filtered using fastp (v0.19.5) (Chen et al., 2018) with parameters “-q 15 -u 40 -n 5 -l 100 --trim_poly_x --detect_adapter_for_pe”.

### Long-read genome assembly, polishing and quality assessment

Porechop (v0.2.3_seqan2.1.1) (https://github.com/rrwick/Porechop) was used to remove adaptors from raw nanopore reads, and Filtlong (v0.2.0) (https://github.com/rrwick/Filtlong) was then used to filter sequences smaller than 1Kb. Three different assemblers: SMARTdenovo (Liu et al., 2021b), Flye (v2.4.2) (Kolmogorov et al., 2019) and WTDBG2 (v2.4.1) (Ruan and Li, 2020), were tested with broccoli nanopore reads that were generated from the first two flow cells. SMARTdenovo was run with parameters “-c 1” to generate consensus sequences and “k −17” to follow developers’ advices for large genomes. Flye was run with parameters “--nano-raw --genome-size 630m”. WTDBG2 was run with preset2 settings “-x ont -g 630m -L 5000 - p 0 -k 15 -AS 2 -s 0.05”. Statistically, SMARTdenovo yielded the most contiguous assembly, with an N50 size of 4.8 Mb and just 472 total contigs (Table S22). The largest contig in this assembly was 19.2 Mb, longer than those for the other two assemblies. Assembly completeness of the three different assemblers was assessed with BUSCO (v3.0.2) (Waterhouse et al., 2018). Among the three assemblers, SMARTdenovo created an assembly with the highest complete BUSCO score (Table S22). Based on these metrics, we selected SMARTdenovo to assemble the five genomes with parameters “-c 1 -k 17”.

Assembled contigs were then polished using nanopore reads for two iterations, followed by Illumina reads for three iterations. For nanopore reads polishing, Minimap2 (v2.18-r1015) (Li, 2018) was used to map raw nanopore reads to raw SMARTdenovo assembly or polished assembly after first round with parameter “-x map-on”. The resulting paf file was submitted to Racon (v1.3.3) for sequence polishing using default parameters (Vaser et al., 2017). For Illumina reads polishing, Illumina paired-end reads were aligned to polished contigs from previous iteration using bwa mem (v0.7.17-r1188) (Li and Durbin, 2009). The resulting bam file was sorted by SAMtools (v1.9) (Li et al., 2009) and then subjected to Pilon (v1.23) (Walker et al., 2014) with default parameters for assembly improvement.

Assembly completeness was assessed by BUSCO (Waterhouse et al., 2018) (embryophyta_odb9 dataset, n=1,440) after each round of polishing. In addition, a variant calling approach was utilized to evaluate the base quality of our assemblies. Illumina short reads were mapped against the raw contig and polished contig sequences using bwa mem (Li and Durbin, 2009), and variations were called with FreeBayes (v1.3.1) (Garrison and Marth, 2012). Only biallelic variants were considered for quality assessment. The total length for each type of variants (SNPs, insertions and deletions) and the total number of bases covered by ≥3X reads were summed with SAMtools depth (Li et al., 2009). Genome-wide quality value (QV) was calculated as 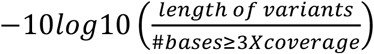 and identity was calculated as 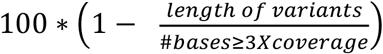 (Jain et al., 2018; Michael et al., 2018).

### DLS optical maps construction and hybrid assembly

High Molecular Weight plant DNA was extracted from fresh tissue of the five morphotypes using the Bionano Prep™ Plant Tissue DNA Isolation Kit. The Direct Label and Stain (DLS) technology, together with Bionano Saphyr platform, were used for generation of optical mapping data. DLS labeling was performed with 750ng DNA using the Direct Labeling and Staining Kit (Bionano Genomics Catalog 80005) following manufacturer’s recommendations. The loading of labeled DNA onto Saphyr chip and running of the Bionano Genomics Saphyr System were all performed according to the Saphyr System User Guide (https://bionanogenomics.com/support-page/saphyr-system/). The generated molecules were *de novo* assembled into genome maps using Bionano Solve Pipeline (version 3.4.1) and Bionano Access (version 1.3). “HybridScaffold” module in Bionano Solve Pipeline was then used to perform hybrid scaffolding between polished contig sequences and Bionano genome maps. As a default parameter, the hybrid scaffolding pipeline didn’t fuse overlapped ONT contigs, which were indicated by the optical maps, but added a 13-bp gap between the two contigs. We checked all 13-bp gaps and aligned both 50-kb flanking regions with BLAT (Kent, 2002). The two flanking contigs were joined if one alignment was detected (Belser et al., 2018). PBJelly (PBSuite_15.8.24) (English et al., 2012) was further used for genome gap filling with the PacBio reads (Table S2).

### TE annotation and gene model prediction

EDTA package (Ou et al., 2019) was used to annotate and classify transposable elements for each assembly. This package selected and combined eight published programs based on benchmarking exercise of a collection of TE annotation programs. Raw candidates from base programs were further filtered to minimize the false discovery rate (Su et al., 2021). Coding sequences from HDEM genome (Belser et al., 2018) were provided for EDTA to remove potential gene-related sequences in the TE library.

Protein-coding gene models were predicted based on repeat-masked assemblies using a strategy that combined *ab initio*, homology-based and transcripts-based predictions. For *ab initio* prediction, Augustus (v3.3.3) (Stanke et al., 2006), SNAP (Korf, 2004) and GlimmerHMM (Majoros et al., 2004) were used to predict gene structures. For homology-based prediction, GeMoMa software (v1.6.3) (Keilwagen et al., 2016) was applied to infer the annotation of protein-coding genes in each of our five assemblies based on protein sequences in previously published genomes, including *Arabidopsis thaliana* (TAIR10, https://www.arabidopsis.org/), *Brassica napus* (Darmor_V8.1), *Brassica napus* (Tapidor_V6.3), *Brassica nigra* (V1.1), *Brassica oleracea* (CAP0212), *Brassica oleracea* (HDEM), *Brassica oleracea* (TO1000), *Brassica rapa* (Chiffu_V3.0) and *Brassica rapa* (Z1). Besides homologous evidences, mRNA-seq data for each morphotype was also incorporated for splice site prediction. GeMoMa was run on each reference genome separately and the resulting gene predictions based on each reference genome were combined. The combined predictions were then filtered to only include complete gene models that were supported by ≥2 reference organisms or mRNA-seq data. For transcripts-based prediction, mRNA-seq reads were assembled into transcripts using two different approaches: *de novo* approach with Trinity (v2.9.1) (Grabherr et al., 2011) and genome-guided approach with Hisat2 (v2.1.0) (Kim et al., 2015) and Stringtie (v2.1.1) (Kovaka et al., 2019). All the transcripts were subject to PASA (v2.4.1) (Haas et al., 2008) for gene model prediction. Finally, EvidenceModeler (v1.1.1) (Haas et al., 2008) was used to combine gene models that were predicted by the three approaches to a weighted consensus gene set.

The protein sequences of predicted gene models were aligned to Swiss-Prot and TrEMBL (Consortium, 2015) databases respectively using diamond (v0.9.32.133) BLASTP with *E* value 1 × 10^-5^. The motifs and domains of protein were predicted by using InterProScan (v5.42-78.0) (Jones et al., 2014) with Pfam, PRINTS, ProSitePatterns, ProSiteProfiles and SMART databases. Gene Ontology (GO) (Ashburner et al., 2000) terms for each gene were extracted from the output of InterProScan. KEGG (Kyoto Encyclopedia of Genes and Genomes) annotation was performed by KAAS (Moriya et al., 2007). TBtools (version 1.09854) (Chen et al., 2020) was used to perform GO enrichment analysis.

### Identification of structural variations

We aligned the other four assemblies to the broccoli genome using minimap2 (v2.18-r1015) (Li, 2018) with parameters “-ax asm5”. The resulting alignments were subject to svim-asm (v1.0.2) (Heller and Vingron, 2020) to call SVs with parameters “haploid --min_sv_size 30 -- max_sv_size 100000”. In addition, we aligned ONT reads from the other four morphotypes to the broccoli genome using NGMLR (v0.2.7) (Sedlazeck et al., 2018) and called SVs using Sniffles (v1.0.12) (Sedlazeck et al., 2018) each with default parameters. To obtain high confidence SV datasets, we then used Jasmine (v1.1.0) (https://github.com/jasmine/jasmine) to merge SVs (insertions and deletions) that were called with different approach and only kept SV that were called by both approach. To identify inversions and translocations, we aligned the other four genomes to the broccoli reference genome using nucmer with parameters “-g 1000 - l 40 -c 90” and filtered the alignments using delta-filter with parameters “-m -i 90 -l 100”. Syri (v1.2) (Goel et al., 2019) was used to identify genomic translocations and inversions based on the alignments.

### Comparative genomics among five *B. oleracea* assemblies

Homologous gene pairs and syntenic relationships between five *B. oleracea* genomes were identified using the MCSCAN toolkit implemented in python (https://github.com/tanghaibao/jcvi/wiki/MCscan-(Python-version)) with default parameters. Microsyntenic dot plots and block depths were generated in python using scripts from MSCAN. The resulting gene pairs were filtered out using a C-score cut-off of 0.99 to obtain 1:1 collinear gene pairs, which were used as input for macrosyntenic analysis with parameters “-- minspan=30 --minsize=30”. To calculate synonymous substitution rates (Ks) for homologous genes among the five assemblies, the identified 1:1 collinear gene pairs were used as input for sequence alignment that was performed using ParaAT_2.0 (Zhang et al., 2012) with parameters “-f axt -m muscle -g”. Ks values were computed based on the alignments using KaKs_Calculator with the method of Nei and Gojobori (Zhang et al., 2006).

### Pan-genome analysis

OrthoFinder (v2.3.12) (Emms and Kelly, 2019) was used to detect orthologous gene clusters based on all protein sequences from nine *B. oleracea* genomes (HDEM, JZS v2.0, Korso, OX-heart and our five genomes) with default parameters. We defined core, dispensable and specific gene clusters as orthologous gene clusters that are present in ≥7 genomes, in 2-6 genomes and in only one genome respectively, and the remaining genes were defined as orphan genes. Pairwise whole-genome sequencing alignments of all possible pairs of the nine genomes were generated using nucmer program in MUMmer4 package (Marçais et al., 2018). The outputs from OrthoFinder and nucmer were used for gene and sequence level pan-genome analyses, respectively, using the approach described in (Jiao and Schneeberger, 2020).

### LTR-RTs analysis

Full-length LTR-RTs were identified by the parallel version of LTR_FINDER (v1.0.7) (Ou and Jiang, 2019; Xu and Wang, 2007) with default parameters, following which LTRharvest (Ellinghaus et al., 2008) was applied with parameters “-minlenltr 100 -maxlenltr 7000 -mintsd 4 -maxtsd 6 -motif TGCA -motifmis 1 -similar 85 -vic 10”. Raw intact LTR-RT candidates that were identified by the two programs were merged. LTR_retriever (v2.8.2) (Ou and Jiang, 2018) was then used to remove false positives and generate non-redundant LTR-RTs. The insertion time of each intact LTR-RT was extracted from the output of LTR_retriever, given the mutation rate of 1.5×10^-8^ mutations per site per year.

### *B. oleracea* subgenome construction and ancestral genome inference

Syntenic gene pairs between nine *B. oleracea* genomes and *A. thaliana* genome were detected using SynOrths (Cheng et al., 2012b). Three subgenomes (the least fractionated (LF), the medium fractionated (MF1) and the most fractionated (MF2) subgenome) of each of the nine *B. oleracea* genomes were constructed using the method reported by Cheng et al (Cheng et al., 2012a). Additionally, a pan-genome based approach was used to infer the ancestral *B. oleracea* genome as reported by Cai et al (Cai et al., 2021). Briefly, we merged all genes in the nine *B. oleracea* genomes that were syntenic to *A. thaliana* genome and ordered these non-redundant genes in the translocation Proto-Calepineae Karyotype (tPCK) (Cheng et al., 2013).

### Phylogenetic tree construction and divergence time estimation

OrthoFinder (Emms and Kelly, 2019) was used to determine single-copy genes between 18 *B. rapa* (Cai et al., 2021), nine *B. oleracea* and *B. nigra* (Perumal et al., 2020) genomes. This resulted in a total of 4,756 single-copy gene families within the 28 genomes. MAFFT (v7.402) (Katoh et al., 2005) was then used to align coding sequences of the single-copy gene families, following which Gblock (v0.91b) (Talavera and Castresana, 2007) was used to extract the conserved sequences among the 28 genomes. IQ-TREE (v1.6.10) (Nguyen et al., 2015) was used to construct Maximum Likelihood tree with the following parameters “-m MFP+ASC -bb 1000 -bnni”. JTT+ASC+R4 was selected as the best model based on the Bayesian Information Criterion (BIC), and 1,000 replicates of ultrafast bootstrapping (UFboot) was used to estimate node support. *B. nigra* was designated the outgroup of the phylogenetic tree. Divergence times were estimated by the program MCMCtree in PAML (paml4.9j) (http://abacus.gene.ucl.ac.uk/software/paml.html) based on the constructed phylogenetic tree. For calibration, we set the divergence time between *B. rapa* and *B. oleracea* at ~4.6 Mya, and between *B. nigra* and the common ancestor of *B. rapa* and *B. oleracea* at ~6.5 Mya (Cheng et al., 2017).

## Acknowledgements

We would like to thank Robin van Velzen (Biosystematics Group, Wageningen University and Research) for helpful discussions.

## Authors’ contributions

GB and CC designed the research. JB performed the experiments. CC and RF analysed data. CC drafted the manuscript. CC and GB revised the manuscript. All authors read and approved the final manuscript.

## Ethics approval and consent to participate

Not applicable

## Consent for publication

Not applicable

