## Supplementary Figures and Tables for "Chromosome-scale genome assemblies of five different *Brassica oleracea* morphotypes provide insights in intraspecific diversification": Supplemenrary figures.pdf

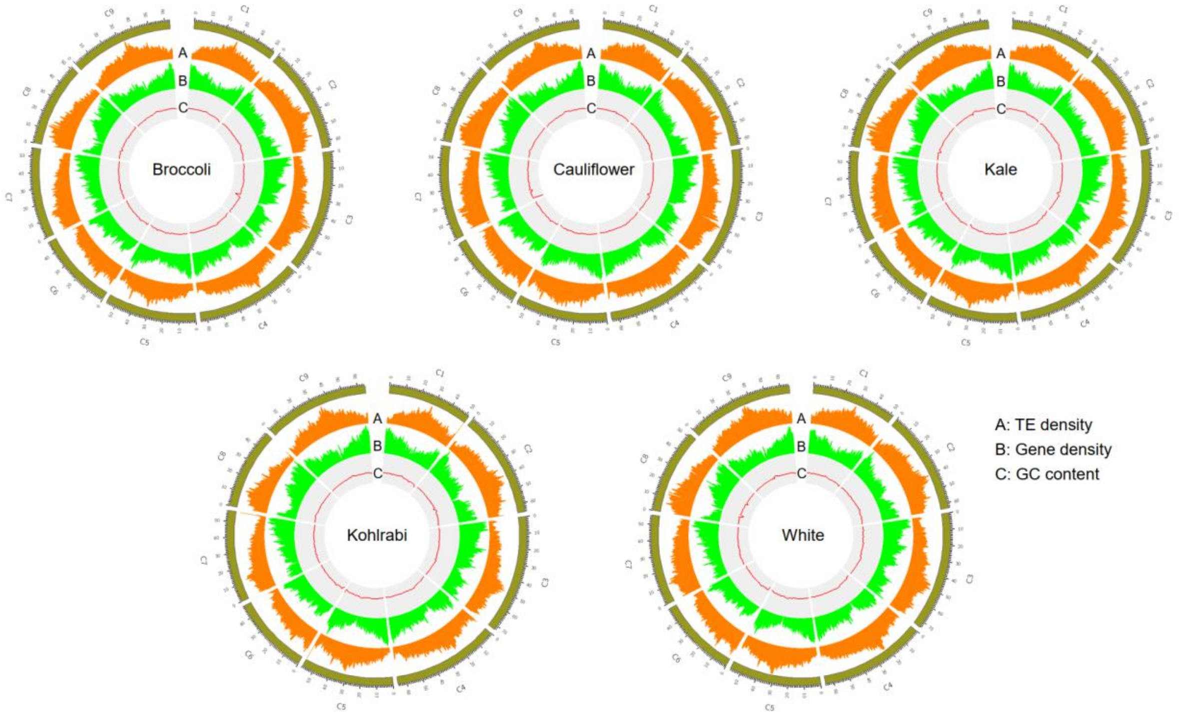


**Fig. S1** Circos plot of genomic landscape in the five *B. oleracea* assemblies. Rings A and B represent transposable-element and gene density in sliding windows of 500Kb with step size of 100Kb. Ring C represents GC content in sliding windows of 2Mb with step size of 1Mb.

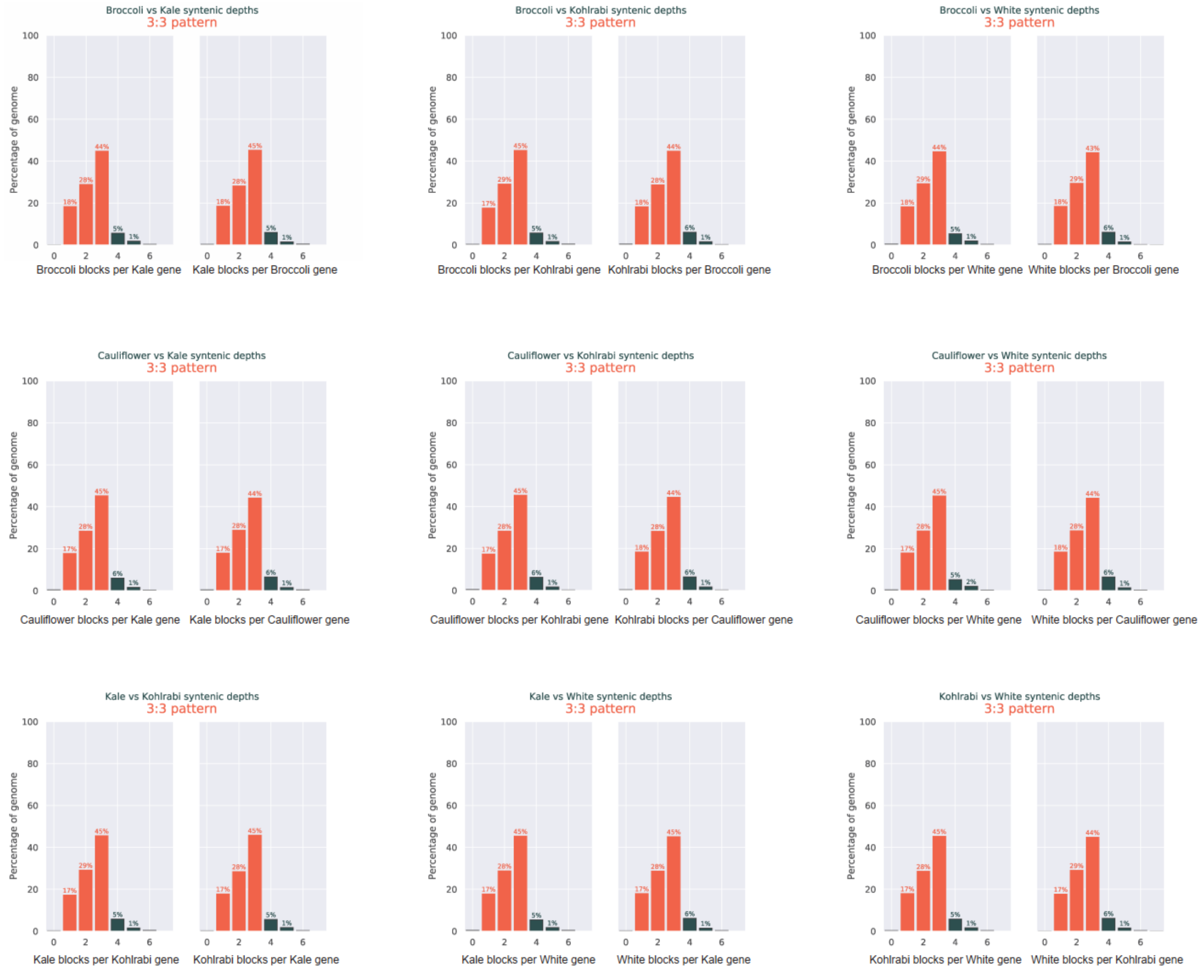


**Fig. S2** Ratio of syntenic depth between each two of the five genomes.

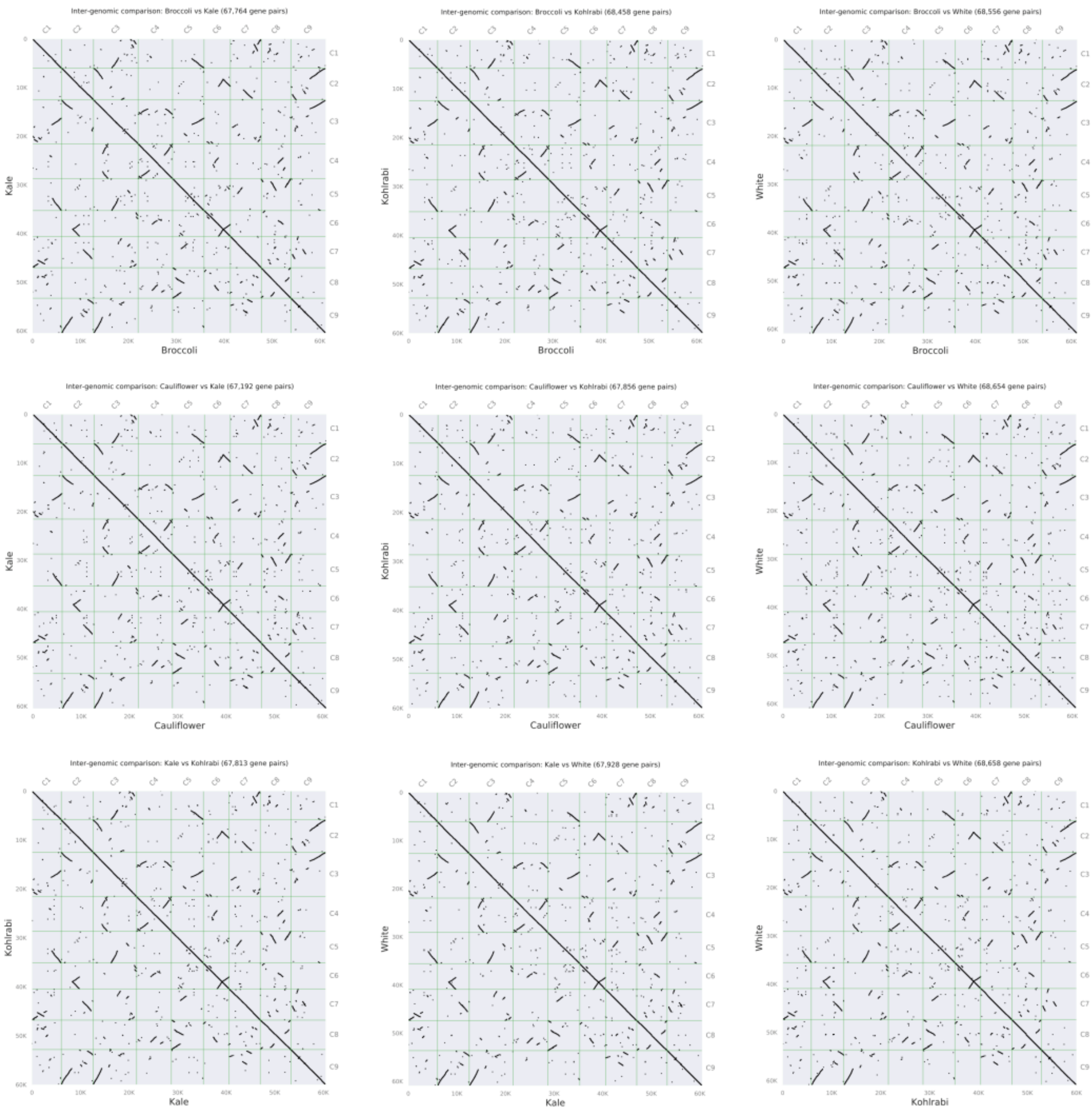


**Fig. S3** Homologous dot plot between each two of the five genomes.

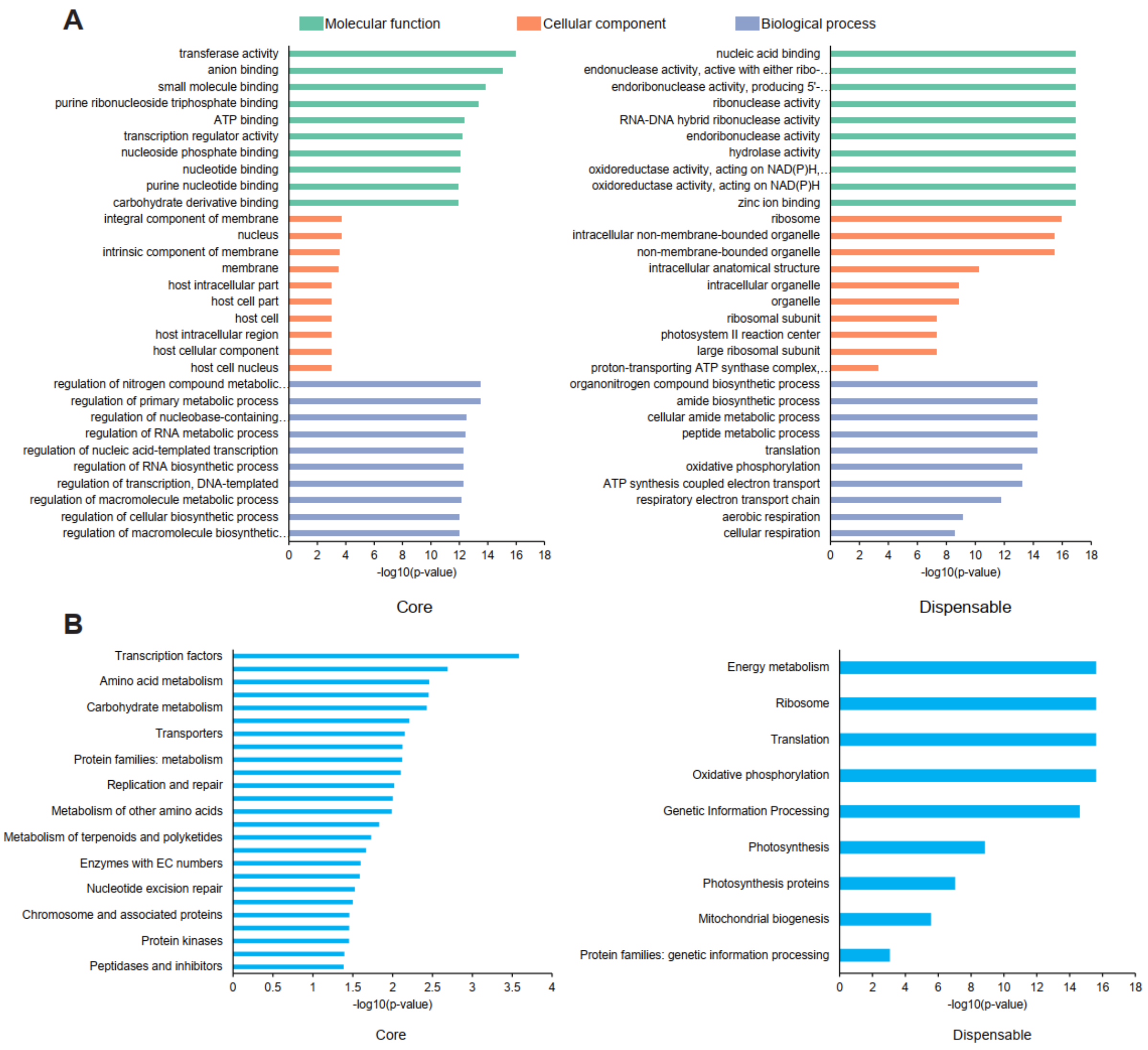


**Fig. S4** Enrichment analysis for core and dispensable gene categories in broccoli genome. (A) Top 10 GO terms enriched in each GO domains (cellular component, biological process, and molecular function) in core and dispensable gene categories. (B) KEGG pathways enriched in core and dispensable gene categories.

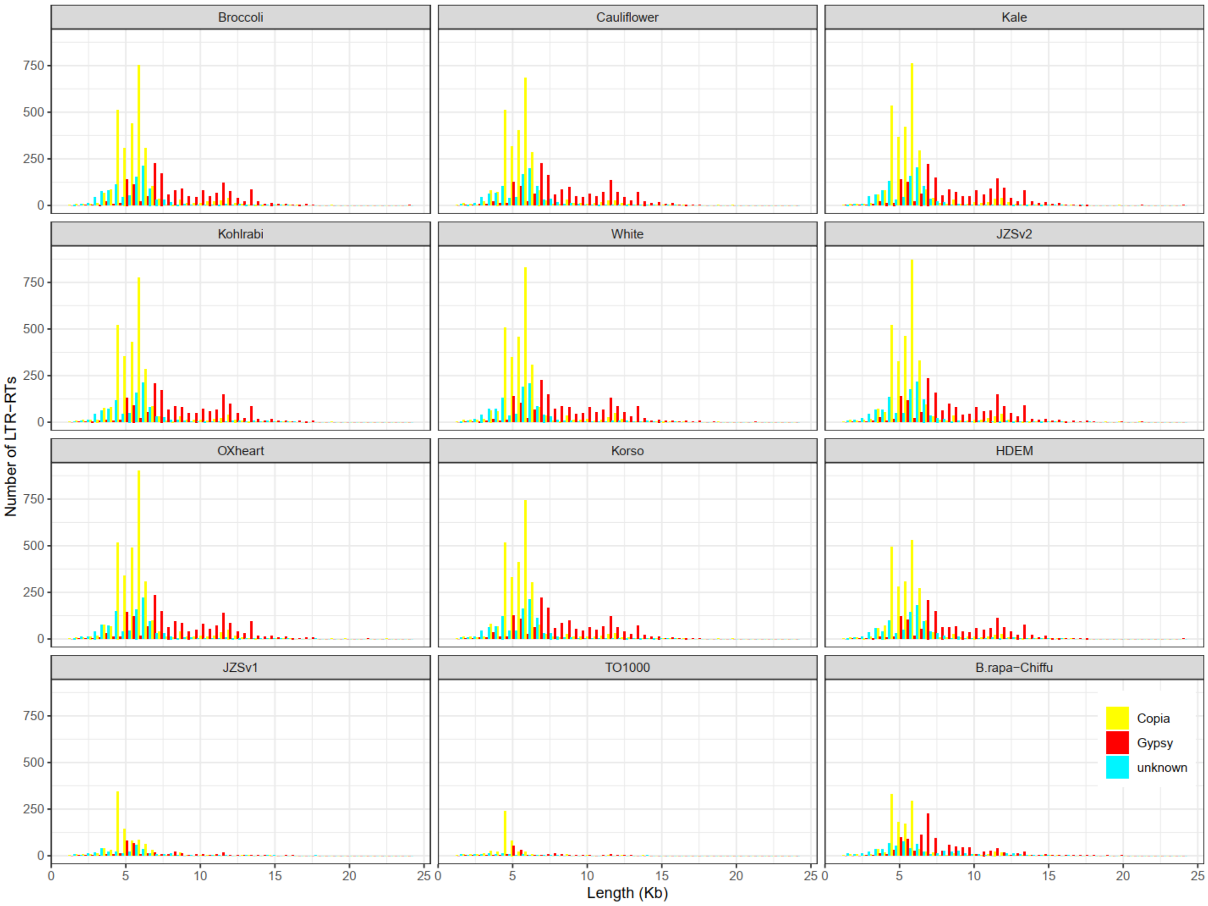


**Fig. S5** Full-length LTR-RTs length distribution in 11 *B. oleracea* and one *B. rapa* genomes.

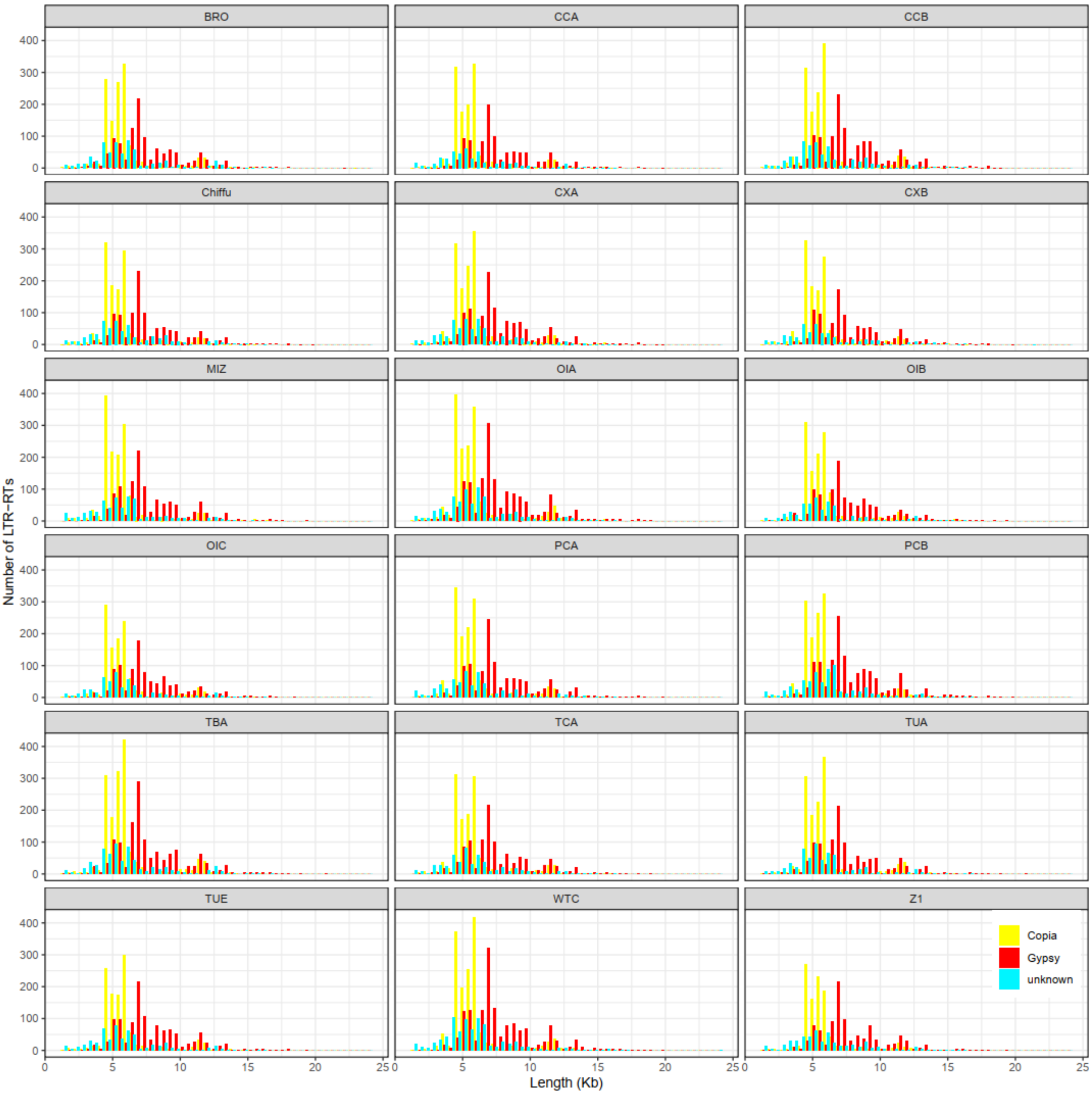


**Fig. S6** Full-length LTR-RTs length distribution in 18 *B. rapa* genomes.

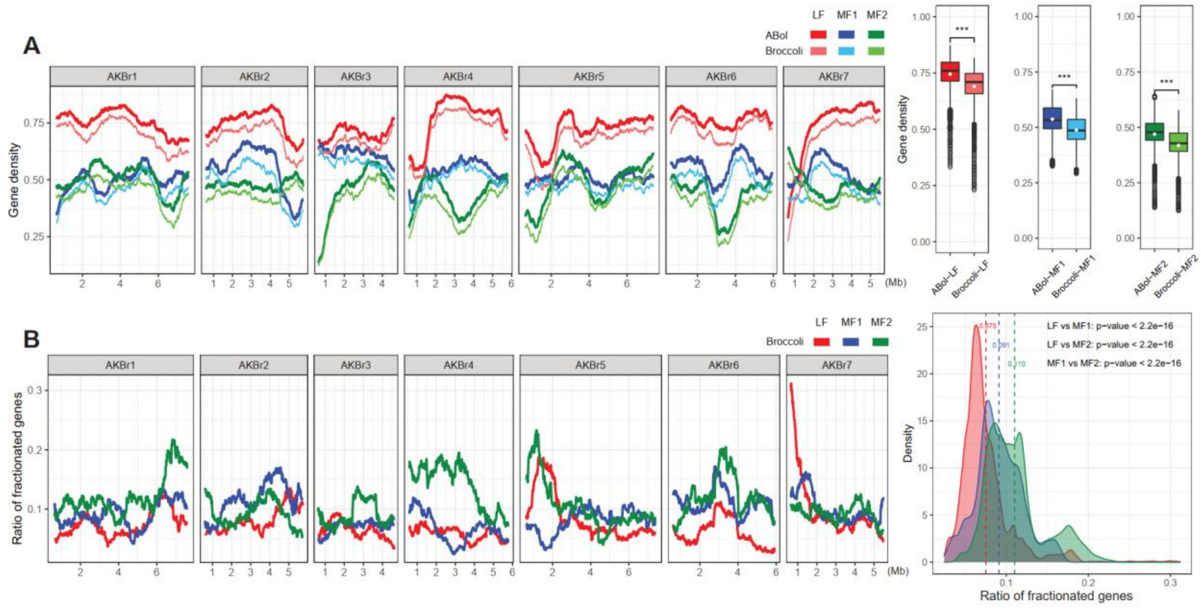


**Fig. S7** Subgenome dominance phenomenon observed in *B. oleracea* during its intraspecific diversification. (A) Gene density distribution on the seven inferred chromosomes of AKBr in the three subgenomes of inferred ancestral genome of *B. oleracea* and broccoli. Broccoli genome was used as a representative to illustrate intraspecies diversification. The figure on the right shows gene density in each window. Two-tailed Student's t-test was performed to compare gene densities between inferred ancestral genome of *B. oleracea* and broccoli for each subgenome. (B) Gene fractionation distribution in the three subgenomes of broccoli. The figure on the right shows the distribution of ratios of fractionated genes to the genes in each window of the inferred ancestral genome, and the dotted line represents the average ratio in each subgenome. 500-gene windows with an increment of two genes was used to calculate gene density and gene fractionation ratio in (A) and (B).

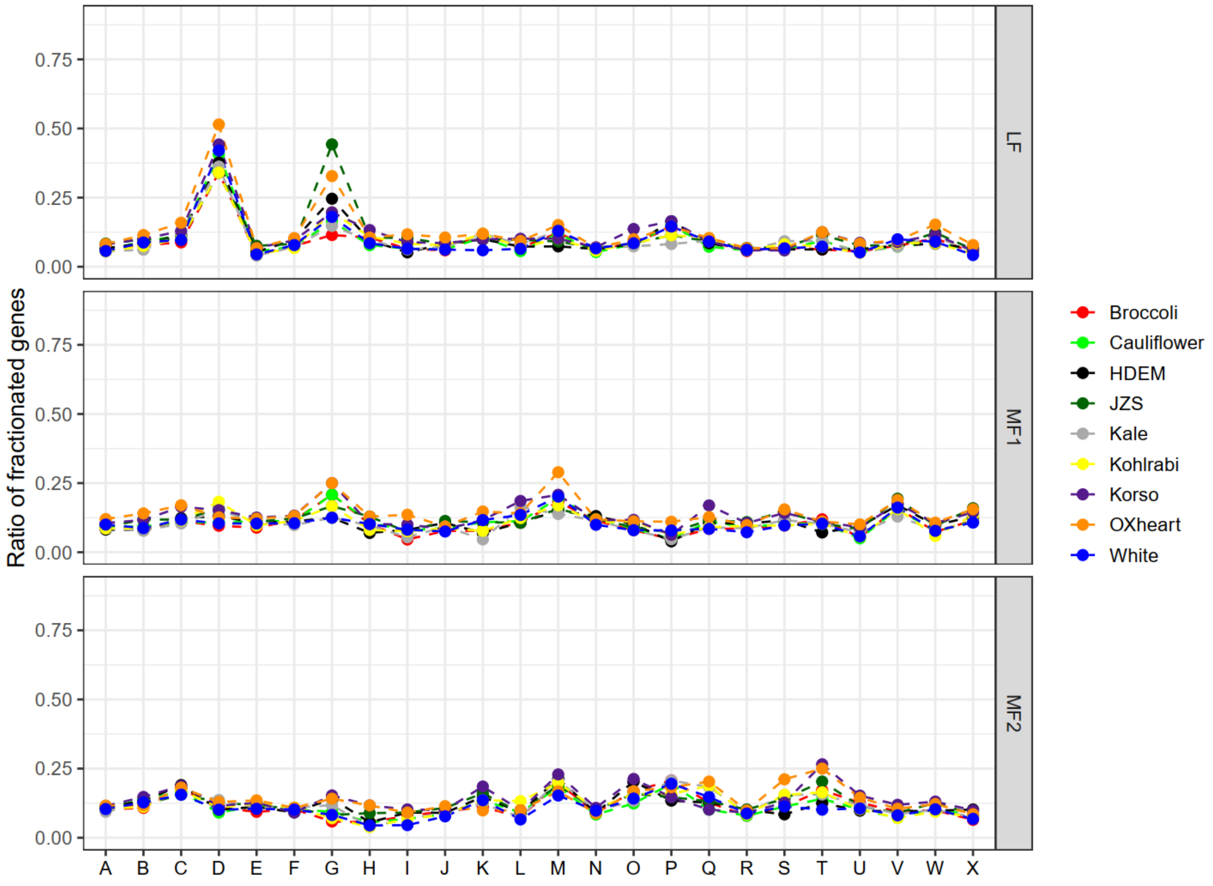


**Fig. S8** Gene fractionation ratios of nine *B. oleracea* accessions in the 24 AK blocks

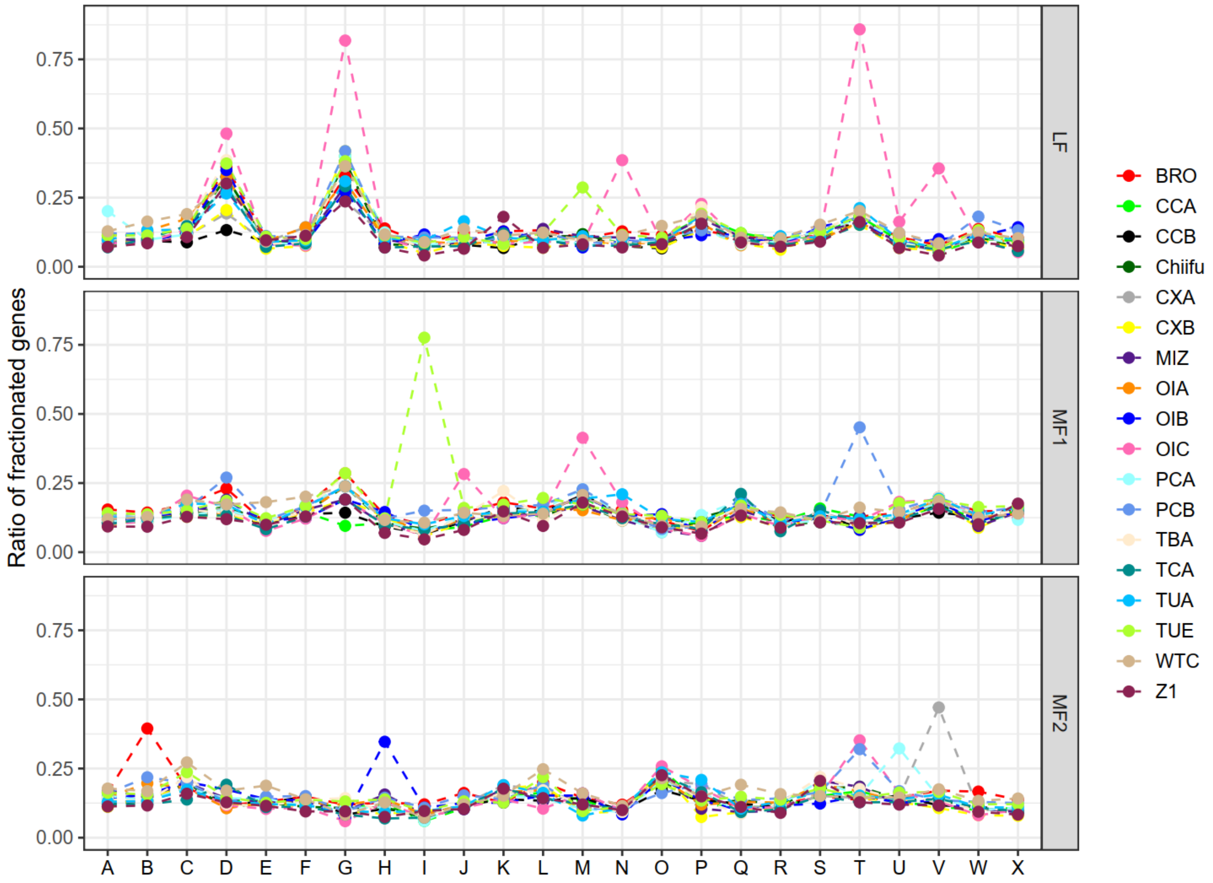


**Fig. S9** Gene fractionation ratios of 18 *B. rapa* accessions in the 24 AK blocks.

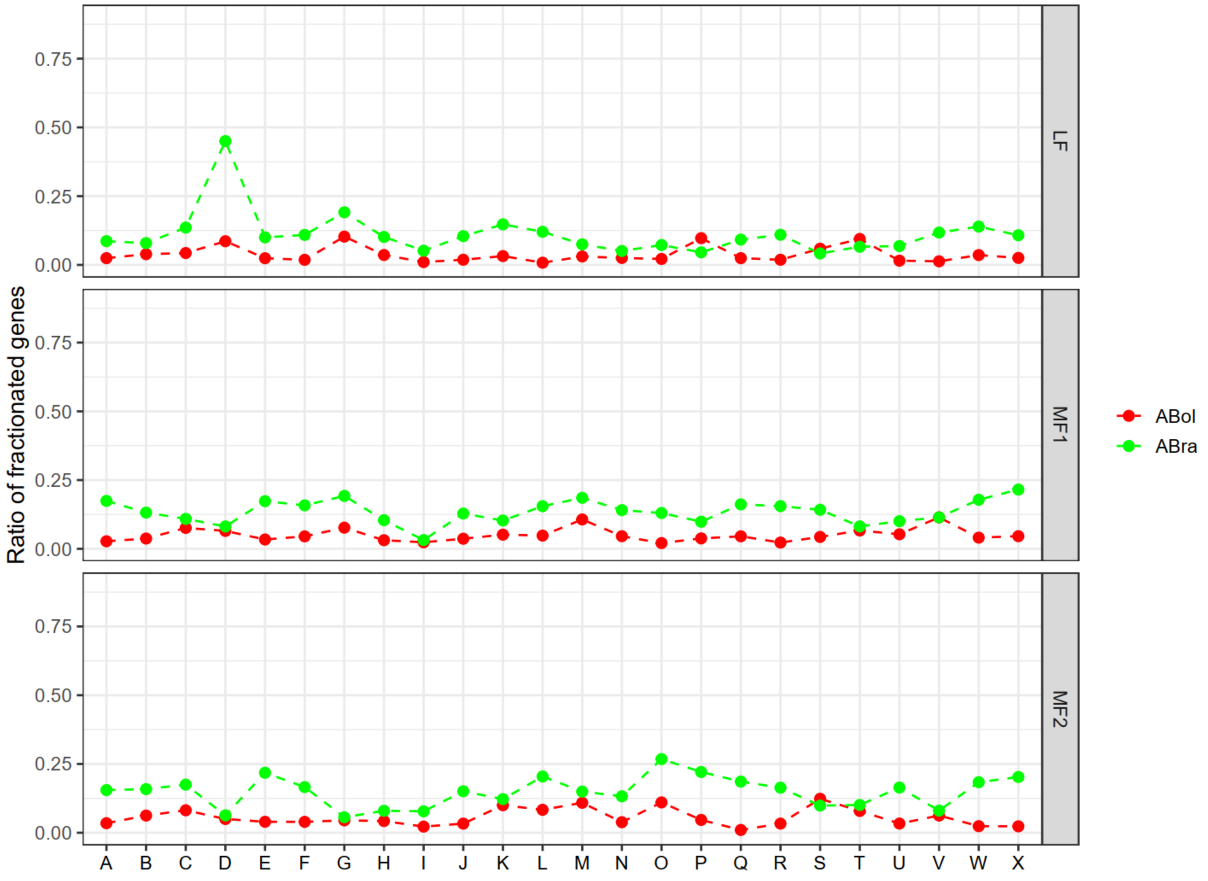


**Fig. S10** Gene fractionation ratios of ancestral genome of *B. oleracea* and *B. rapa* in the 24 AK blocks.

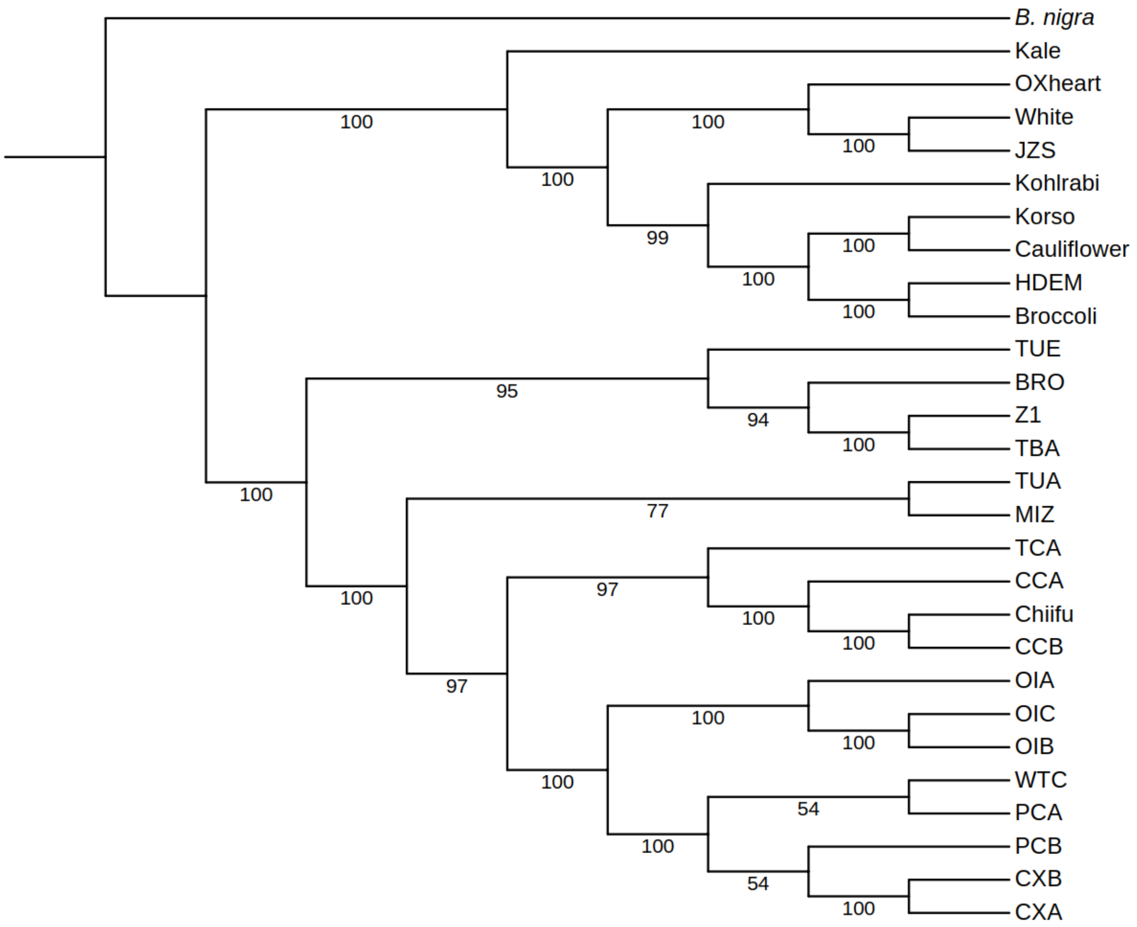


**Fig. S11** Phylogenetic relationships of nine *B. oleracea* and 18 *B. rapa* accessions using *B. nigra* as an outgroup. Numbers below each node show the bootstrap values.

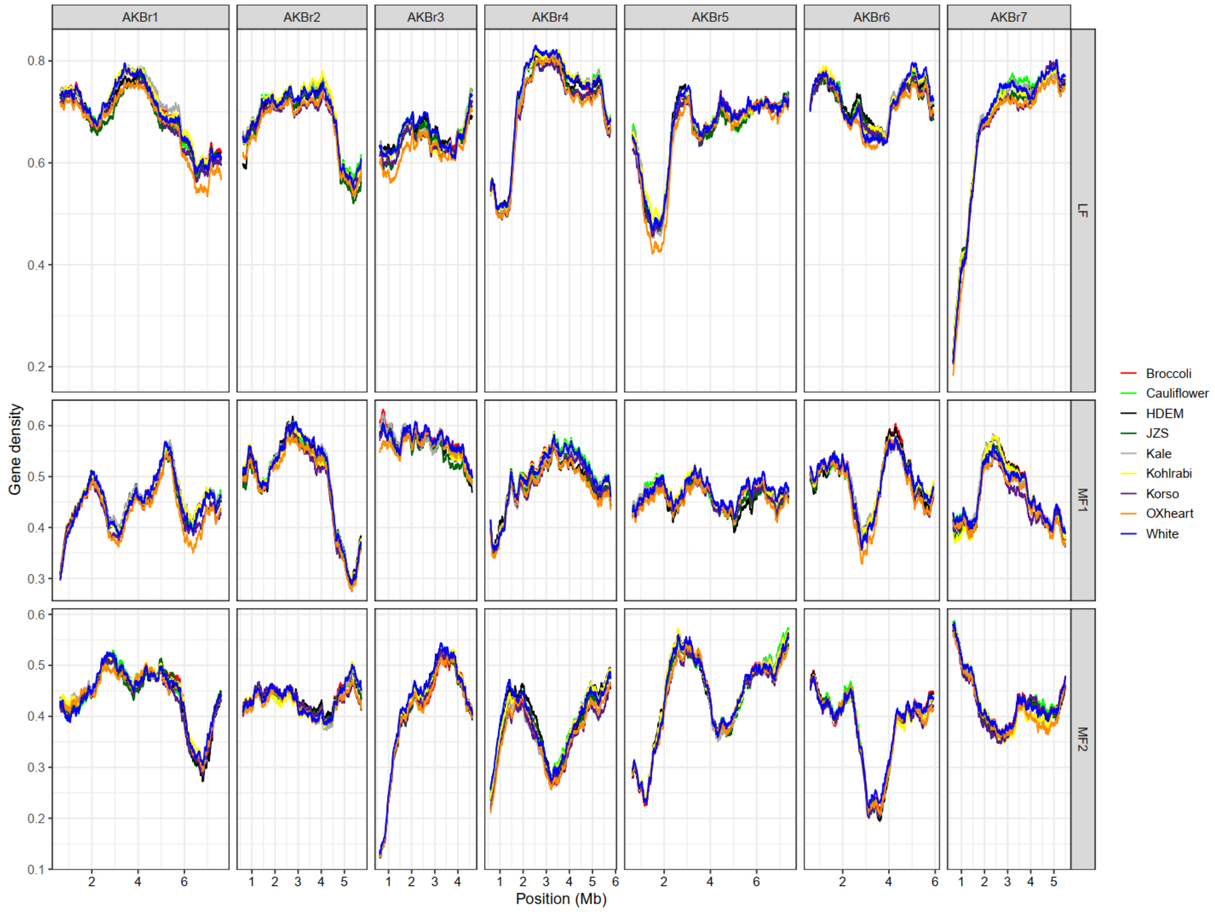


**Fig. S12** Gene density distribution on the seven inferred chromosomes of AKBr in the three subgenomes of nine *B. oleracea* accessions. 500-gene windows with an increment of two genes was used to calculate gene density.

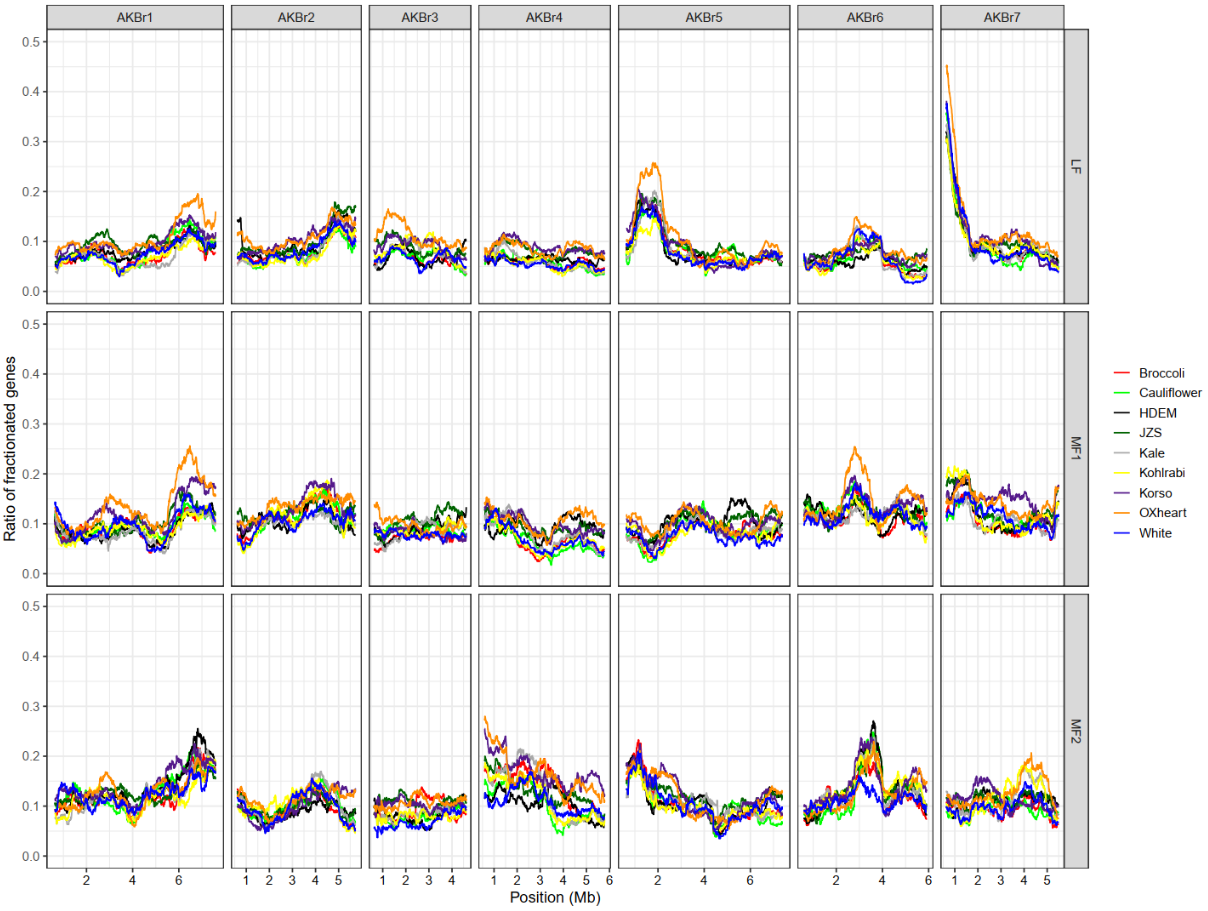


**Fig. S13** Gene fractionation distribution on the seven inferred chromosomes of AKBr in the three subgenomes of nine *B. oleracea* genomes. 500-gene windows with an increment of two genes was used to calculate gene fractionation ratio.

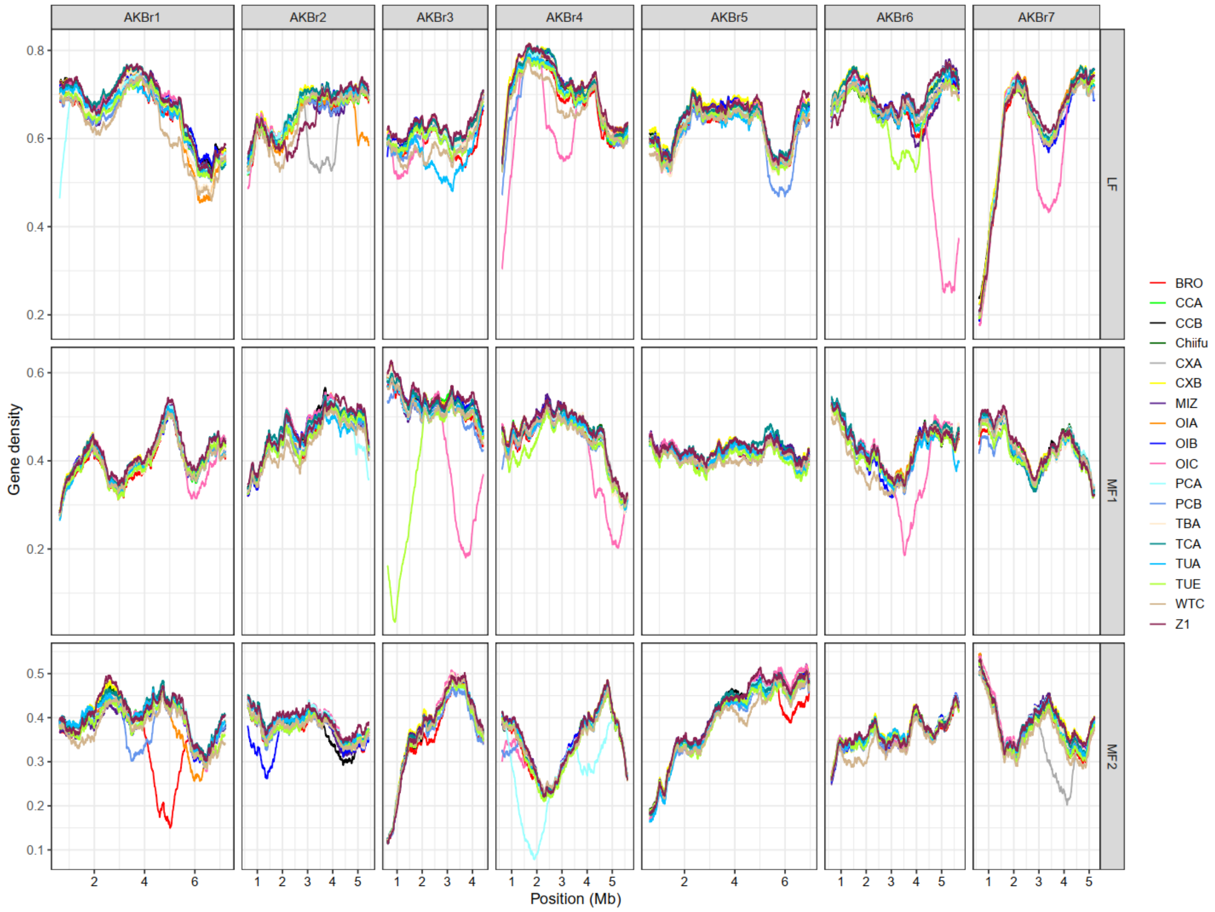


**Fig. S14** Gene density distribution on the seven inferred chromosomes of AKBr in the three subgenomes of 18 *B. rapa* accessions. 500-gene windows with an increment of two genes was used to calculate gene density.

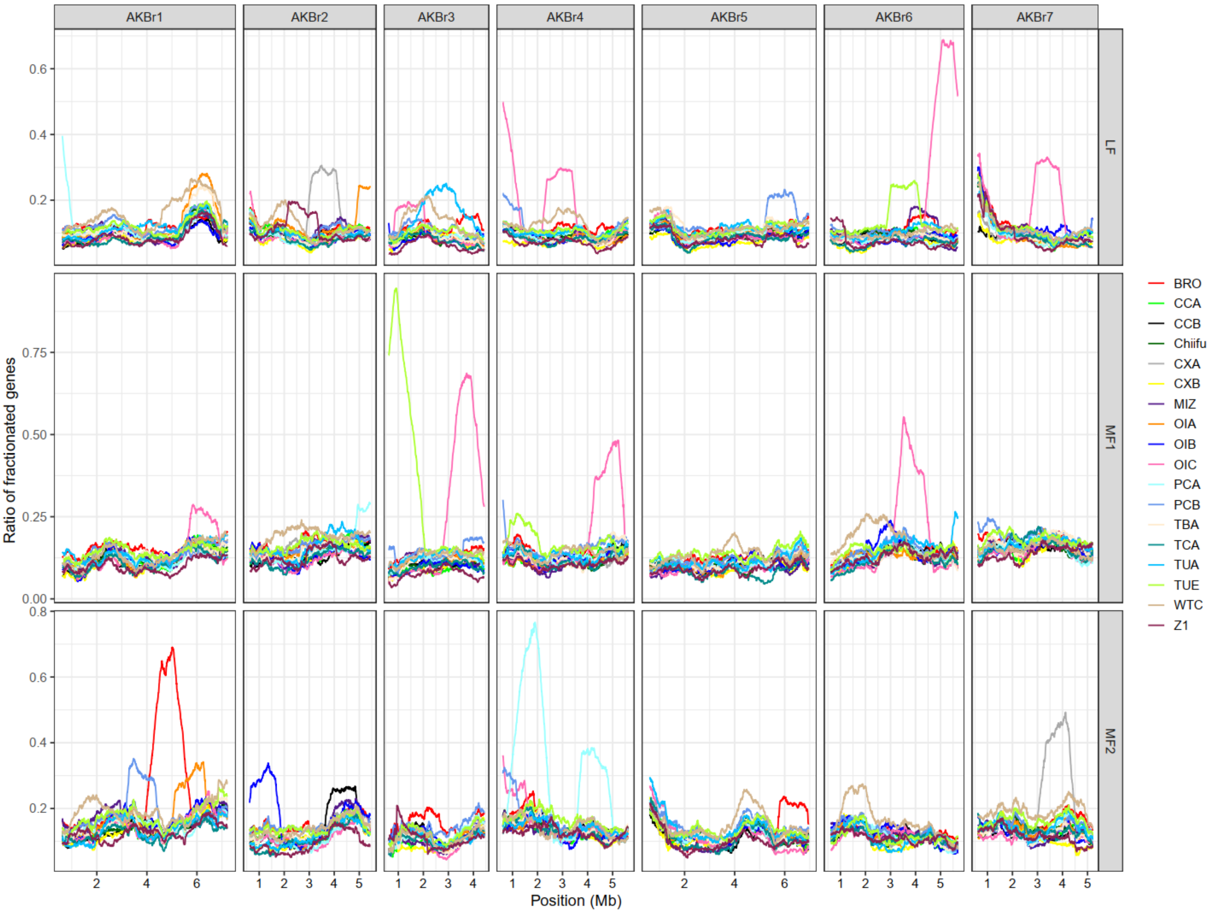


**Fig. S15** Gene fractionation distribution on the seven inferred chromosomes of AKBr in the three subgenomes of 18 *B. rapa* genomes. 500-gene windows with an increment of two genes was used to calculate gene fractionation ratio.

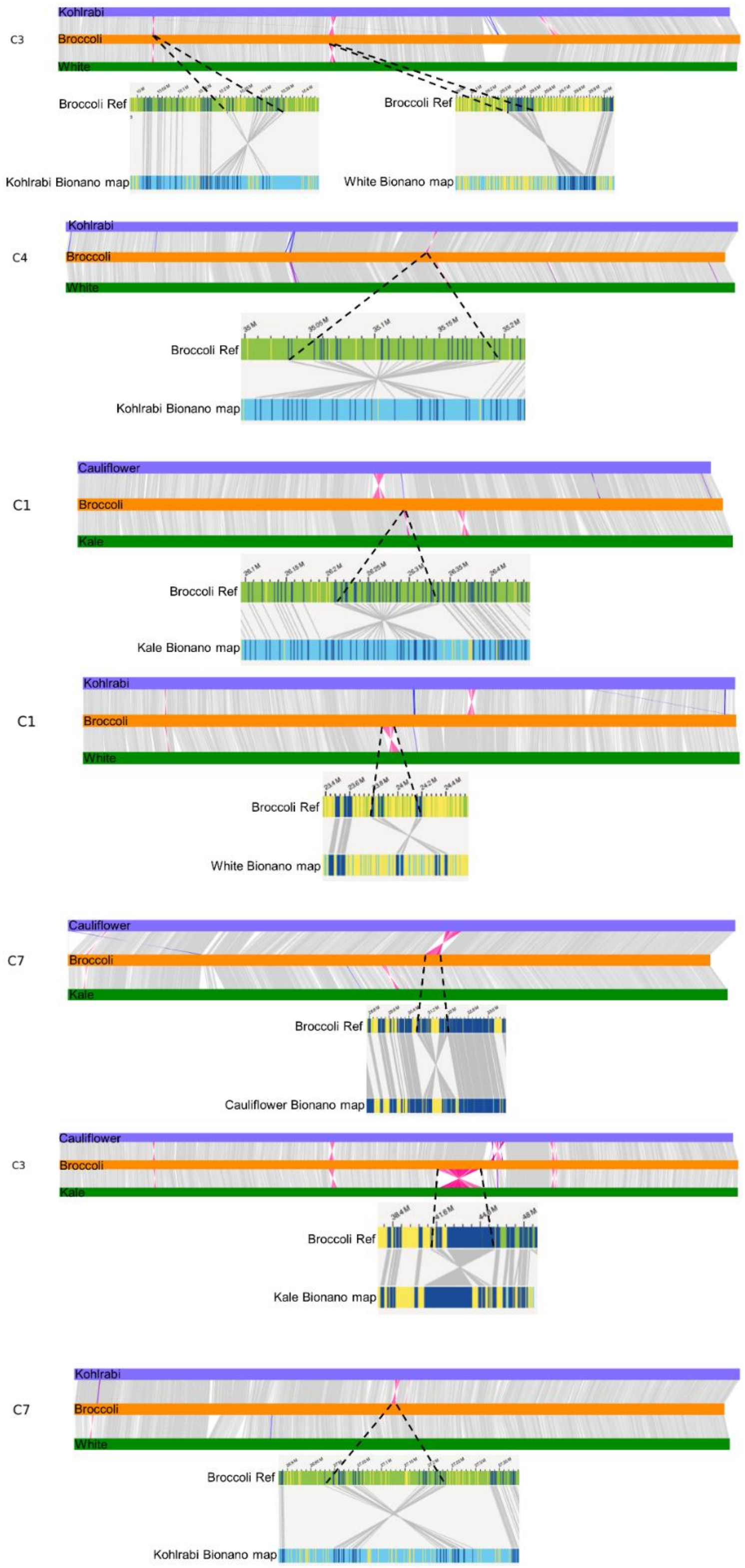


**Fig. S16** Eight large inversions that were supported by Bionano maps.

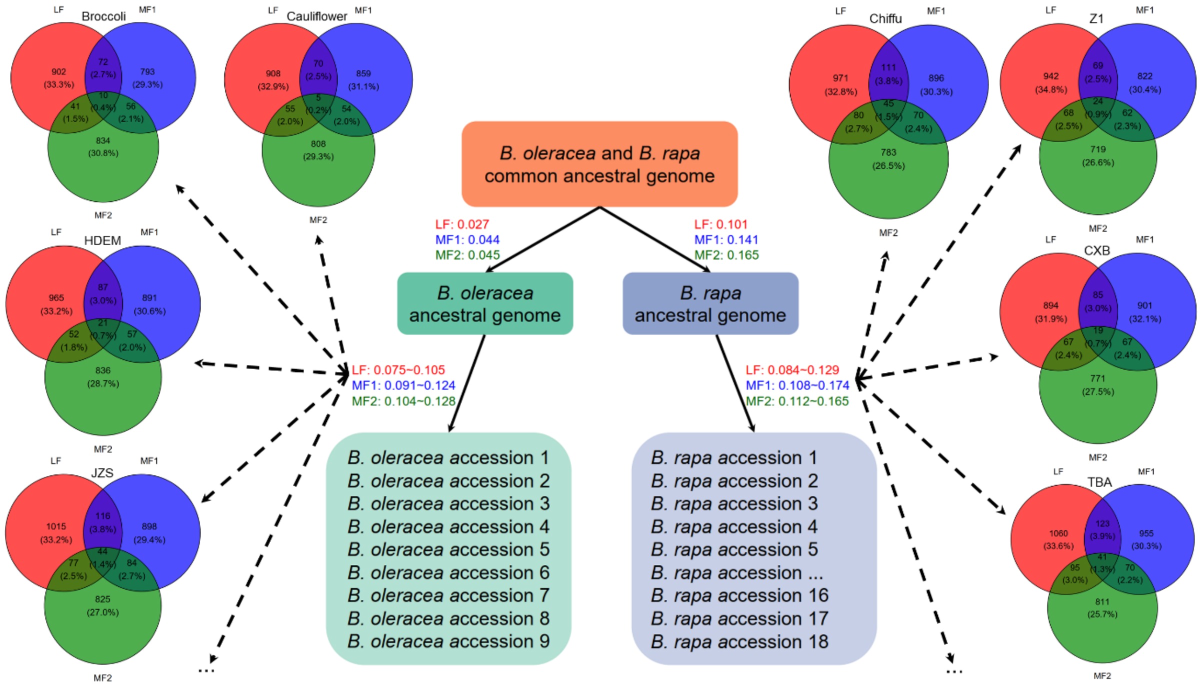


**Fig. S17** One-copy gene loss bias during intraspecific diversification of *B. oleracea* and *B. rapa*.

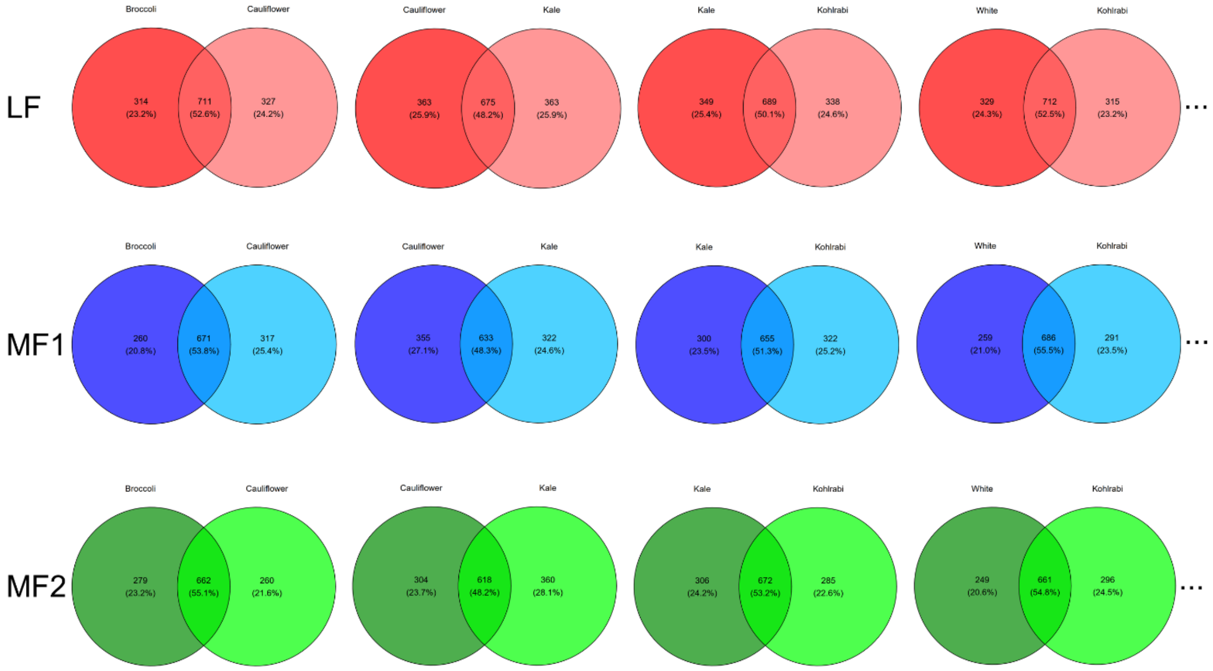


**Fig. S18** Examples of common and shared gene loss between two extant *B. oleracea* genomes in the three subgenomes.

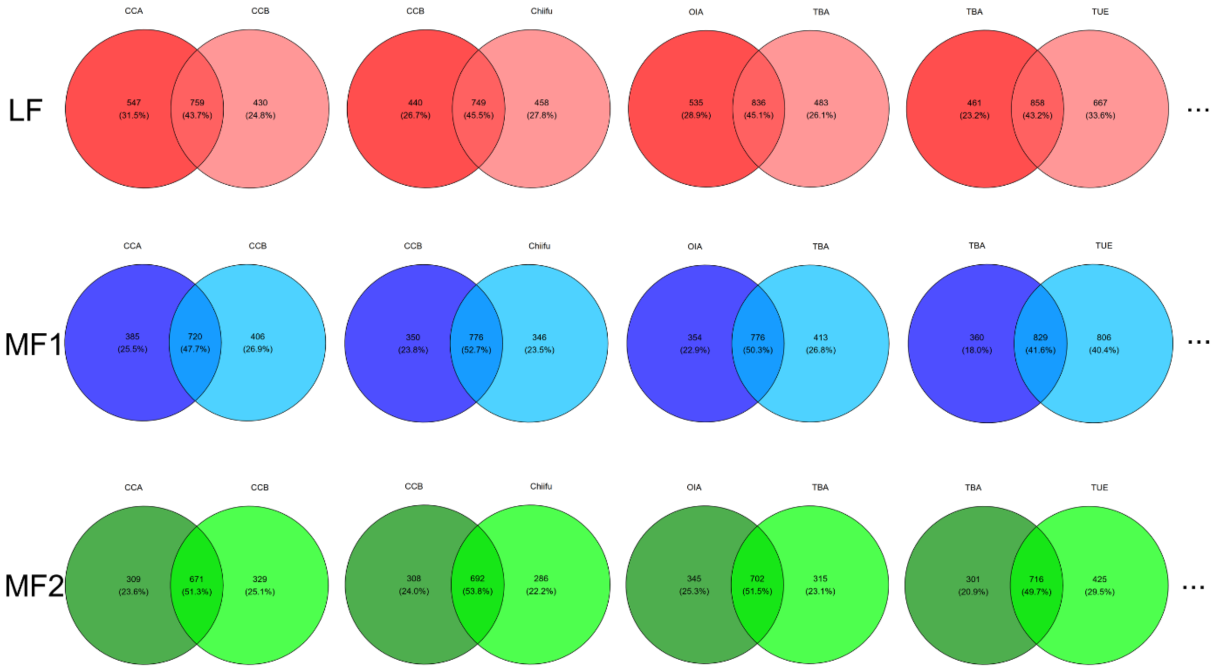


**Fig. S19** Examples of common and shared gene loss between two extant *B. rapa* genomes in the three subgenomes.

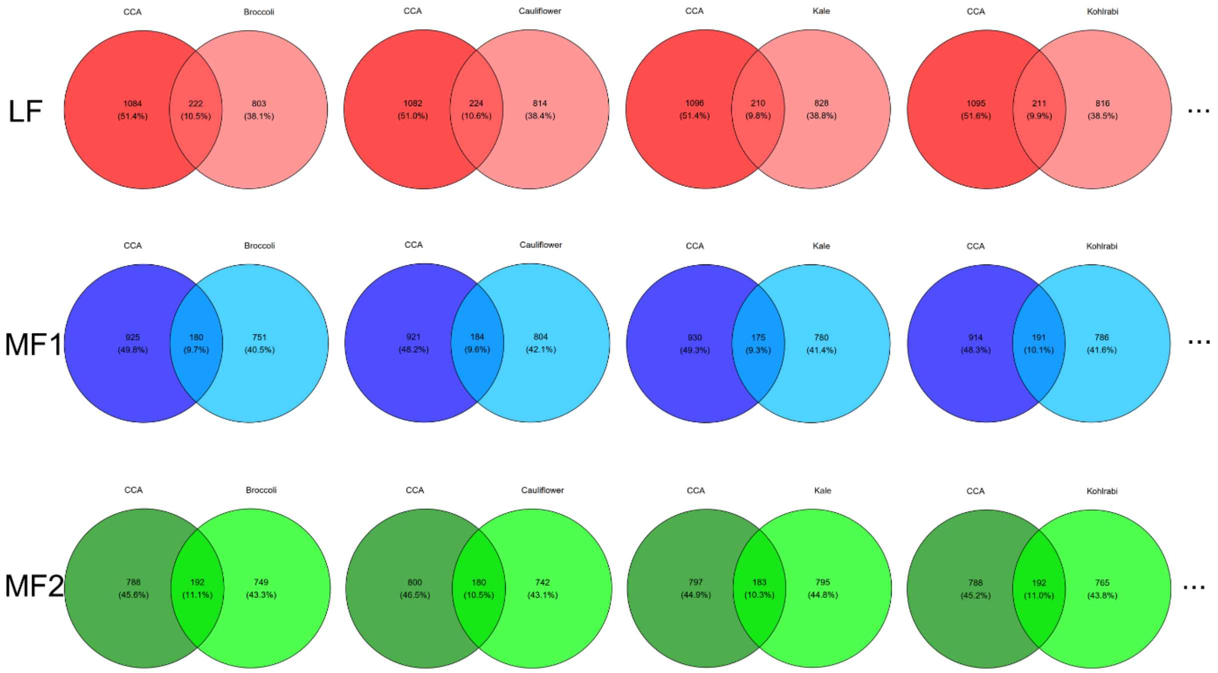


**Fig. S20** Examples of common and shared gene loss between extant *B. rapa* and *B. oleracea* genomes in the three subgenomes.
